# A highly sensitive cell-based luciferase assay for high-throughput automated screening of SARS-CoV-2 nsp5/3CLpro inhibitors

**DOI:** 10.1101/2021.12.18.473303

**Authors:** KY Chen, T Krischuns, L Ortega Varga, E Harigua-Souiai, S Paisant, A Zettor, J Chiaravalli, D Courtney, A O’Brien, SC Baker, C Isel, F Agou, Y Jacob, A Blondel, N Naffakh

## Abstract

Effective drugs against SARS-CoV-2 are urgently needed to treat severe cases of infection and for prophylactic use. The main viral protease (nsp5 or 3CLpro) represents an attractive and possibly broad-spectrum target for drug development as it is essential to the virus life cycle and highly conserved among betacoronaviruses. Sensitive and efficient high-throughput screening methods are key for drug discovery. Here we report the development of a gain-of-signal, highly sensitive cell-based luciferase assay to monitor SARS-CoV-2 nsp5 activity and show that it is suitable for high-throughput screening of compounds in a 384-well format. A benefit of miniaturisation and automation is that screening can be performed in parallel on a wild-type and a catalytically inactive nsp5, which improves the selectivity of the assay. We performed molecular docking-based screening on a set of 14,468 compounds from an in-house chemical database, selected 359 candidate nsp5 inhibitors and tested them experimentally. We identified four molecules, including the broad-spectrum antiviral merimepodib/VX-497, which show anti-nsp5 activity and inhibit SARS-CoV-2 replication in A549-ACE2 cells with IC_50_ values in the 4-21 µM range. The here described assay will allow the screening of large-scale compound libraries for SARS-CoV-2 nsp5 inhibitors. Moreover, we provide evidence that this assay can be adapted to other coronaviruses and viruses which rely on a viral protease.

## 1. Introduction

In January 2020, a novel human coronavirus named severe acute respiratory syndrome coronavirus 2 (SARS-CoV-2) was identified as the causative agent of the COVID-19 disease. SARS-CoV-2 has spread globally, causing a pandemic that is still on-going and has to date infected over 237 million people and caused over 4.6 million deaths worldwide [1]. Vaccines started to be deployed early 2021 and more than 6.4 billion doses have been administered to date [1]. However, until complete vaccine coverage will be reached and with the recent emergence of variants of concern, there is a critical need for prophylactic and therapeutic antiviral drugs. Despite huge efforts, only few treatment have been approved or authorized under emergency use authorization [2], among which the glucocorticoid dexamethasone [3] and the IL6 receptor blocker tocilizumab [4], which show a benefit in severe COVID-19 cases. Other treatments include the REGN-COV2 antibody cocktail [5], whose high cost precludes large-scale administration, and the ribonucleotide analog remdesivir whose efficacy is limited [6]. The identification of additional anti-SARS-CoV-2 drugs therefore remains a high priority.

SARS-CoV-2 is an enveloped virus with a single-stranded, positive-strand RNA genome. It belongs to the *Coronaviridae* family, which includes other zoonotic coronaviruses (SARS-CoV-1 and the Middle-East Respiratory Syndrome Coronavirus or MERS-CoV) as well as human seasonal coronaviruses that cause common colds. The large open reading frame ORF1ab at the 5’ end of the SARS-CoV-2 genome is translated into viral polyproteins that are cleaved by two viral proteases into at least 16 non-structural proteins (nsps) [7]. The main protease is a chymotrypsin-like cysteine-dependent protease called nsp5 (or 3CLpro or Mpro), and the second protease is a papain-like protease domain within nsp3 called PLpro (for a review see [8]). Both proteases represent attractive targets for drug development as they are essential to the progression of the viral life cycle, and their active sites are highly conserved among betacoronaviruses. Nsp5 is a dimer composed of two identical monomers of 306 residues, while the PLpro region of nsp3 is 317 residues long and composed of 4 subdomains, including an ubiquitin-like domain. Nsp5 is considered to be a particularly attractive target because it cleaves the viral polyprotein 1ab at 11 major cleavage sites and is required to produce the precursors of the viral replication complex (nsp12-nsp7-nsp8), and because its cleavage specificity is distinct from human proteases. To date, there are more than 250 crystal structures available for the SARS-CoV-2 nsp5, including co-crystal structures with small molecules, which facilitates structure-based design of new antivirals [9-14]. Early on, the lopinavir/ritonavir protease inhibitor cocktail that prevents HIV replication was proposed as an antiviral agent against SARS-CoV-2, however early clinical trials did not indicate benefits in patients with severe COVID-19 [15]. Boceprivir, a protease inhibitor used to treat patients with Hepatitis C virus (genotype 1) chronic infection, showed only a moderate activity against SARS-CoV-2 nsp5 [16, 17] and was not tested in COVID-19 clinical trials. The covalent peptidomimetics PF-00835231 initially developed by the Pfizer company against SARS-CoV-1 and MERS-CoV and its prodrug PF-07304814 are also active against SARS-CoV-2 [18]. Pfizer recently announced that its novel oral antiviral Paxlovid, a combination of PF-07321332 [19, 20] and ritonavir, reduced the risk of hospitalization or death by nearly 90% compared to placebo in interim analysis of a Phase 2/3 trial in high-risk adults with COVID-19 (NCT04960202 and NCT05011513). Only a few other studies with protease inhibitors are currently registered on Clinical.Trials.gov, notably a phase II study on Masitinib (NCT05047783).

Paxlovid is currently being reviewed by the FDA and EMA and may obtain an Emergency Use Authorization in the near future, which would represent a new and major tool for pandemic management. Although the structure of the active site of nsp5 is highly conserved among coronaviruses, and its sequence shows very few variations among the > 1.9 millions SARS-CoV-2 full-genome available to date [21], the emergence of viruses with drug-resistant mutations cannot be excluded. Therefore, sensitive and efficient high-throughput screening methods are key for the discovery of additional novel protease inhibitors. *In vitro* biochemical assays, which assess the protease activity of purified SARS-CoV-2 nsp5 are available [10, 13, 22-25]. However, *in vitro* assays need to be complemented with cell-based assays as they do not take into account compound membrane permeability, toxicity and bioavailability in cells. Cell-based reporter assays that can be performed in a Biosafety Level 1 (BSL1) setting have been developed [26-31] but their suitability for high-throughput screening has not been demonstrated. Here we report the development of a rapid and highly sensitive cell-based luciferase assay to monitor SARS-CoV-2 nsp5 activity. We provide evidence that the reporter assay is suitable for high-throughput automated screening of large libraries of compounds and report the identification of 4 lead compounds that inhibit protease activity and SARS-CoV-2 replication in cell culture.

## 2. Material and methods

*Antibodies and Virtual screening for anti-nsp5 compounds are described in the Supplementary Methods*

### Cells and virus

HEK-293T cells (purchased from ATCC, # CRL-11268) and A549 cells stably expressing the human ACE2 receptor (A549-ACE2, kindly provided by O. Schwartz, Institut Pasteur) were grown in complete Dulbecco’s modified Eagle’s medium (Gibco) supplemented with 10% fetal bovine serum (FBS). Blasticidin was added to the A549-ACE2 medium at a concentration of 10 µg/mL to maintain the expression of ACE2. The BetaCoV/France/IDF0372/2020 virus was kindly provided by S. van der Werf (Institut Pasteur).

### Plasmids

The pcDNA3.1 expression vectors for a SARS-CoV-2 polyprotein corresponding to the non-structural proteins 4, 5 and the N-terminal part of nsp6 (designated CoV-2-nsp4-5-6 WT), and its catalytically inactive counterpart in which cysteine 145 (with numbering starting at the first residue of nsp5) was changed to alanine (designated CoV-2-nsp4-5-6 C145A) are described in [31]. The mutated cleavage site between nsp5 and 6 was changed to the wild-type sequence by site-directed mutagenesis. These plasmids were used as templates to amplify the sequence encoding nsp5 (WT or C145A), and the resulting amplicons were subcloned into the pcDNA3.1 plasmid. The pLEX expression vectors for SARS-CoV-2, IBV and hCoV-229E nsp5 are described in [26] and were kindly provided by N. Heaton (Duke University). The Hepatitis C Virus (HCV) NS3/4A expression vector was generated by subcloning a synthetic gene which was codon optimized for expression in human cells (Genscript) into the pCI plasmid. Site-directed mutagenesis was performed to generate a catalytically inactive mutant in which S139 (with numbering starting at the first residue of NS3) was changed to alanine (designated HCV-NS3/4A S139A). To generate the reverse-Nanoluciferase reporter plasmid Rev-Nluc-CoV, a synthetic codon-optimized nucleotide sequence encoding a permuted Nanoluciferase with the NGSVRLQSSLK linker between the N- and the C-terminal domains was cloned into the pLV-CMV-eGFP lentiviral vector (Duke vector core, Duke University) in place of the eGFP insert. The other Rev-Nluc reporter plasmids were produced in two steps. First, a synthetic codon-optimized nucleotide sequence encoding a permuted Nanoluciferase with two BsmBI sites between the N- and the C-terminal domain was cloned into pLV-CMV-eGFP. Second, double-stranded oligonucleotides encoding alternative nsp5, Tobacco Etch Virus (TEV) or HCV-NS3/4A protease cleavage sites were cloned in frame using the BsmBI sites. All constructs were verified by sequencing. Primers used for cloning and mutagenesis are available upon request.

### Viral protease activity assays

HEK-293T cells were seeded in 96-well white opaque plates (Greiner Bio-One, 3×10^4^ cells per well) one day before being transfected with 5 ng of the Rev-Nluc-CoV reporter plasmid and 95 ng of the pcDNA3.1-nsp4-5-6 WT, pcDNA-3.1-nsp4-5-6 C145A, pLEX-nsp5 or an empty control pCI plasmid, using polyethyleneimine (PEI-max, #24765-1 Polysciences Inc). To measure NS3/4A activity, 5 ng of the Rev-Nluc-HCV reporter plasmid was co-transfected with 145 ng of the pCI-NS3/4A WT or pCI-NS3/4A S139A, using the same protocol. Nanoluciferase activity was measured at 24 or 48 hours post-transfection (hpt) using the Nano-Glo® Luciferase Assay System (#1120, Promega) and a Berthold Centro XS3 luminometer (integration time 1s/well using the MikroWin® software). When indicated, cells were treated at 8 hpt with increasing concentrations of the inhibitors GC376 (Sigma-Aldrich), Calpain Inhibitor XII (Cayman Chemicals), Mpro 13b (Bio-Techne) and Boceprevir (Sigma-Aldrich), with a constant DMSO concentration of 0.1%, for ∼14 h. Nanoluciferase activity was measured at 24 hpt.

In the 384-well plate setting, ∼10^7^ HEK-293T cells were transfected in 50 cm^2^ dishes with 1 µg of the Rev-Nluc-CoV plasmid and 15 µg of the nsp4-5-6 WT or nsp4-5-6 C145A expression plasmids. Increasing amounts of the tested compounds, corresponding to final concentrations of 0.1 to 50 µM with a constant DMSO concentration of 0.5% were distributed in 384-well white opaque plates using an Echo 555 Liquid Handler (Labcyte). Transfected HEK-293T cells were trypsinized at 6 hpt and distributed in the 384-well plates (2×10^4^ cells per well in 50 µL). The luciferase activity was measured at 24 hpt as described above. A Tecan-Fluent robotic workstation was used to dispense cells and luciferase substrate. Each 384-well plate included 14 negative control wells (mean DMSO-treated signal = DMSO) and 14 positive-control wells (mean GC376 at 50 µM signal = GC). The compound luciferase signals (compound-treated signal = C) measured in nsp4-5-6 WT expressing cells were normalized as follows: (C - DMSO) / (GC – DMSO) x 100. The luciferase signals measured in nsp4-5-6 C145A expressing cells were normalized as follow: (C / DMSO) x 100.

Additional information about tested compounds (MolPort Compound number, supplier, purity) is provided in the Supplemental Material and Methods section.

### Antiviral activity and cell viability assays

A549-ACE2 cells seeded in 384-well plates (1.5×10^3^ cells per well) were incubated with the compound of interest at the indicated concentration in DMEM-2% FBS 2 h prior to infection. The media was replaced with the SARS-CoV-2 inoculum (MOI = 0.1 PFU/cell) and incubated for 1 h at 37 °C. The inoculum was removed and replaced with drug-containing media. At 72 hpi, the cell culture supernatant was collected, heat inactivated at 80 °C for 20 min, and used for RT-qPCR analysis. The Luna Universal One-Step RT-qPCR Kit (New England Biolabs) was used, with SARS-CoV-2 specific primers targeting the N gene region (5’-TAATCAGACAAGGAACTGATTA-3’ and 5’-CGAAGGTGTGACTTCCATG-3’) and with the following cycling conditions: 55 °C for 10 min, 95 °C for 1 min, and 40 cycles of 95 °C for 10 sec, followed by 60 °C for 1 min, in an Applied Biosystems QuantStudio 6 thermocycler. The percentage of viral growth inhibition was calculated as the ratio of the sample mean Ct to the positive control (remdesivir 1 µM) mean Ct, after subtracting the negative control (DMSO 0.5 %) mean Ct. Cell viability assays were performed in drug-treated cells using the CellTiter-Glo assay according to the manufacturer’s instructions (Promega) and a Berthold Centro XS LB960 luminometer. The percentage of cytotoxicity was calculated relative to untreated cells (0% cytotoxicity) and cells treated with 20 µM camptothecin (100% cytotoxicity).

## 3. Results

### 3.1. Development of a reverse-Nanoluciferase (Rev-Nluc) reporter to monitor nsp5 activity

To develop a Nanoluciferase-based reporter for SARS-CoV-2 nsp5 activity, we built on our experience in engineering a split Nanoluciferase for protein complementation assays [32]. We designed a reverse-Nanoluciferase (Rev-Nluc-CoV, **Figure 1A**) in which the two independent structural Nanoluciferase domains corresponding to amino acids 2-66 and 66-171 are permuted and separated by a flexible loop including the coronavirus nsp5 consensus cleavage site VRLQ/S [33]. As a control, a TEV protease cleavage site or a sequence corresponding to two BsmBI restriction sites was inserted into the Rev-Nluc (Rev-Nluc-TEV and Rev-Nluc-control, respectively). To determine the efficiency of nsp5-mediated Rev-Nluc cleavage, the Rev-Nluc-CoV plasmid was co-transfected with a plasmid encoding nsp4, nsp5 and the amino-terminal part of nsp6 [31]. Previous studies have shown that nsp5 is released upon autocatalytic processing of the coronavirus nsp4-5-6 polyprotein [31, 33, 34]. As a control, a catalytically inactive nsp5 mutant (C145A) was used [34].

**Figure 1.**
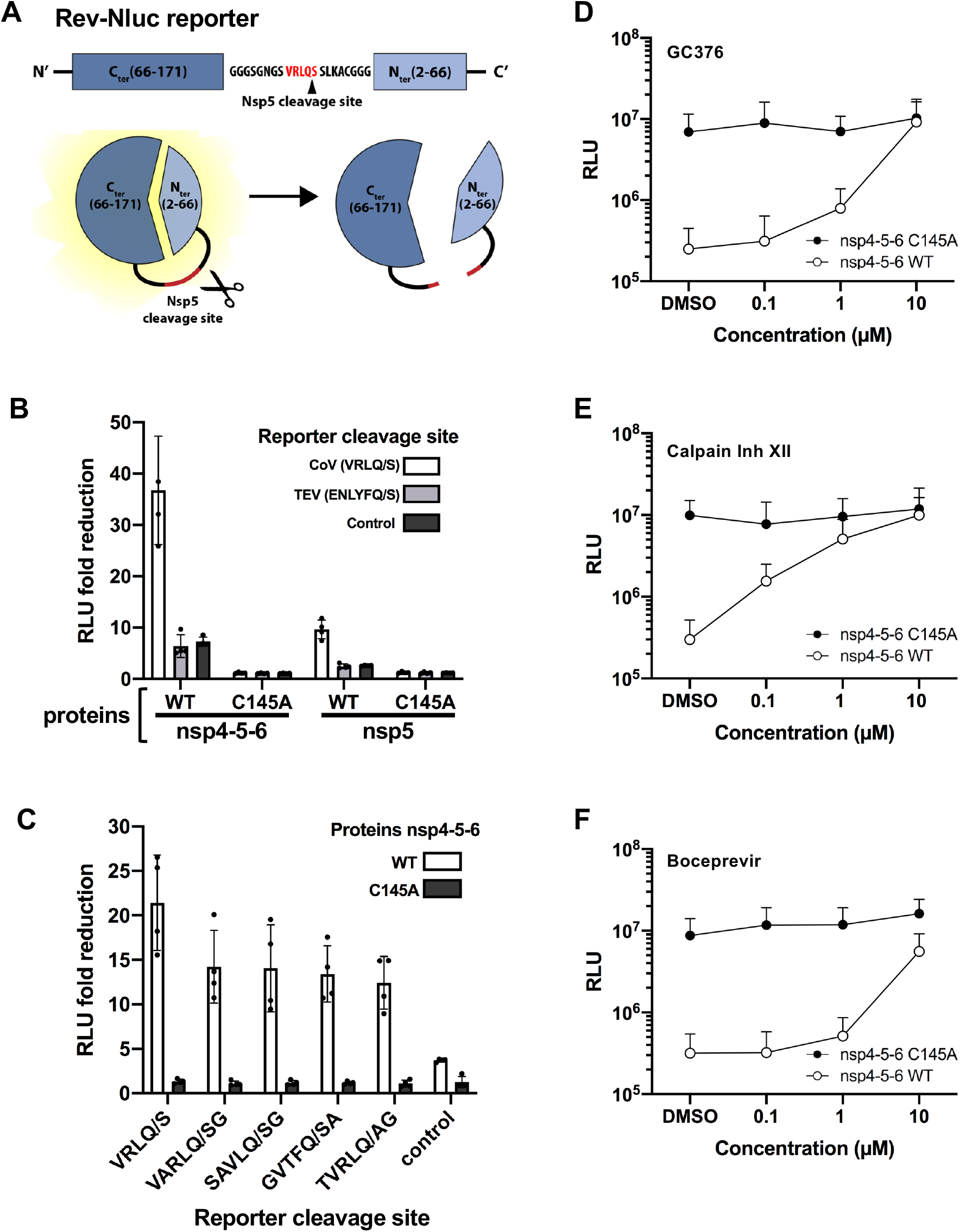
Development of a Nanoluciferase-based reporter to monitor nsp5 activity. **A**. Schematic representation of the Rev-Nluc-CoV based assay for nsp5/3CLpro activity. The Rev-Nluc-CoV reporter was designed so that the N-terminal and C-terminal domains of the Nanoluciferase are permuted and separated by a flexible loop including the coronavirus nsp5 consensus cleavage site (VRLQ/S). Reporter cleavage by nsp5 reduces Nanoluciferase activity. **B**. Luciferase signals measured at 48 hpt in HEK-293T cells transfected with a Rev-Nluc reporter (CoV cleavage site, TEV cleavage site or a control linker sequence) and nsp4-5-6 (WT or C145A) or nsp5 (WT or C145A). The graph shows the Relative Light Unit (RLU) fold reduction compared to cells transfected with an empty control plasmid instead of the nsp4-5-6 or nsp5 plasmid (mean ± SD of four independent experiments performed in technical triplicates). **C**. Luciferase signals measured at 24 hpt in HEK-293T cells transfected with a Rev-Nluc reporter (with different nsp5 cleavage sites or a control linker sequence), and nsp4-5-6 (WT or C145A). The graph shows the RLU fold reduction compared to cells transfected with an empty control plasmid instead of the nsp4-5-6 plasmid (mean ± SD of four independent experiments performed in technical triplicates). **D-F**. The inhibition of SARS-CoV-2 nsp5 by GC376 (D), Calpain Inhibitor XII (E) and Boceprevir (F) was assessed using the Rev-Nluc-based assay. The assay was performed as described in the Methods section. The RLU are shown as the mean ± SD of three independent experiments performed in technical triplicates. Open symbols: nsp4-5-6 WT; closed symbols: nsp4-5-6 C145A.

A luciferase signal was measured in cells co-transfected with the Rev-Nluc-CoV and a control empty plasmid, demonstrating that the permuted Nanoluciferase domains form an active enzyme. Compared to the empty vector control, cells co-transfected with the SARS-CoV-2 nsp4-5-6 WT plasmid showed a ∼30-fold reduction in luciferase signal at 48 hpt, whereas Nanoluciferase signals in cells co-transfected with the SARS-CoV-2 nsp4-5-6 C145A remained unchanged (**Figure 1B**, white bars, and **Supplemental Figure 1**). When the Rev-Nluc-TEV and Rev-Nluc-control plasmids were used instead of the Rev-Nluc-CoV reporter plasmid, cells co-transfected with nsp4-5-6 WT showed a ∼5-fold decrease in luciferase signal (**Figure 1B**, light and dark grey bars, respectively). These data indicate that upon autocatalytic processing of the SARS-CoV-2 nsp4-5-6 polyprotein, the released nsp5 restricts the production of an active Nanoluciferase, and it does so through two distinct mechanisms: 1) cleavage of the canonical nsp5 site VRLQ/S in the Rev-Nluc and 2) reduction of Nanoluciferase activity independent of the canonical cleavage site, possibly caused by nsp5-mediated cleavage of Rev-Nluc at other sites or indirectly by cleavage of cellular proteins. The cleavage of the reporter is strictly dependent on the protease activity of nsp5, as demonstrated by the nsp5 C145A mutant. When nsp5 was expressed alone instead of the nsp4-5-6 polyprotein, the observed fold-changes were lower (**Figure 1B**).

An orthogonal assay was used to confirm that the reduction in luciferase signal was due to nsp5-mediated cleavage of the reporter. The Rev-Nluc-CoV reporter was fused to the 3xFLAG tag at its N-terminal end and cleavage was assessed by western-blot using an anti-FLAG antibody. As shown in **Supplemental Figure 2**, the full-length Rev-Nluc-CoV protein could no longer be detected upon co-expression of the SARS-CoV-2 nsp4-5-6 WT, while its levels remained unchanged upon expression of the C145A mutant. Notably, the FLAG-tagged cleavage product with an expected size of 19 kD (compared to 30 kD for the uncleaved Rev-Nluc) could not be detected either, suggesting that it is degraded upon cleavage.

### Comparison of Rev-Nluc reporters with different nsp5 cleavage sites

In an attempt to optimize the Rev-Nluc reporter, we replaced the initial coronavirus nsp5 consensus VRLQ/S cleavage site by the VARLQ/SG sequence that was identified as an optimal SARS-CoV nsp5 cleavage site [35], or by three nsp5 cleavage sites in the SARS-CoV-2 ORF1a at the nsp4-5 (SAVLQ/SG), nsp5-6 (GVTFQ/SA) and nsp9-10 (TVRLQ/AG) junctions. With all reporter plasmids, a decrease in luciferase signal was measured in cells expressing SARS-CoV-2 nsp4-5-6 WT (**Figure 1C**, white bars) compared to the empty vector, but not in cells expressing the mutant SARS-CoV-2 nsp4-5-6 C145A (**Figure 1C**, dark grey bars). The highest fold change was observed with the Rev-Nluc-CoV plasmid (VRLQ/S), which was used in the following experiments in combination with SARS-CoV-2 nsp4-5-6 WT or C145A.

### Effect of nsp5 inhibition on Rev-Nluc activity

The fold-change in Nanoluciferase signal measured between the WT and C145A mutant is dependent on nsp5 enzymatic activity and can therefore be used as a proxy for *in vivo* nsp5 activity. To investigate whether the Rev-Nluc reporter could be used to screen for inhibitors of nsp5, we tested the commercially available GC376 compound that was reported to inhibit SARS-CoV-2 nsp5 activity *in vitro* [17, 22] as well as in cell-based assays [26, 27] and to inhibit SARS-CoV-2 replication in Vero cells [23]. GC376 dose-dependently reduced nsp5 activity, as seen by a gradual increase of Nanoluciferase activity measured in the presence of SARS-CoV-2 nsp4-5-6 WT (**Figure 1D**, open symbols), while the luciferase signal measured in the presence of catalytically inactive mutant remained unchanged (**Figure 1D**, closed symbols). Full inhibition of nsp5 was achieved at 10 µM GC376 and the estimated IC_50_ was in the micromolar range, in agreement with previous findings in cell-based assays [26, 36, 37]. The inhibitory effects of Calpain Inhibitor XII and Boceprevir as determined by the Rev-Nluc assay were lower compared to GC376 (**Figures 1E** and **1F**), consistent with the reported *in vitro* IC_50_ values (0.45 and 4.13 µM respectively, compared to 0.03 µM for GC376 in [17]).

### Extended use of the Rev-Nluc assay

We asked whether the Rev-Nluc-CoV reporter can be used to monitor the activity of nsp5 proteins from distantly related coronaviruses. To this end, the Rev-Nluc-CoV plasmid was co-transfected with nsp5 expression vectors derived from SARS-CoV-2 (*Betacoronavirus*), the human seasonal coronavirus 229E (hCoV-229E, *Alphacoronavirus*) or the avian infectious bronchitis virus (IBV, *Gammacoronavirus*). Compared to the empty vector control, a 50- to 100-fold reduction was observed in the presence of SARS-CoV-2, hCoV-229E and IBV nsp5 (**Figure 2A**). These findings are consistent with data obtained with an nsp5 FlipGFP reporter [26], and show that the Rev-Nluc assay can be adapted to other coronaviruses.

**Figure 2.**
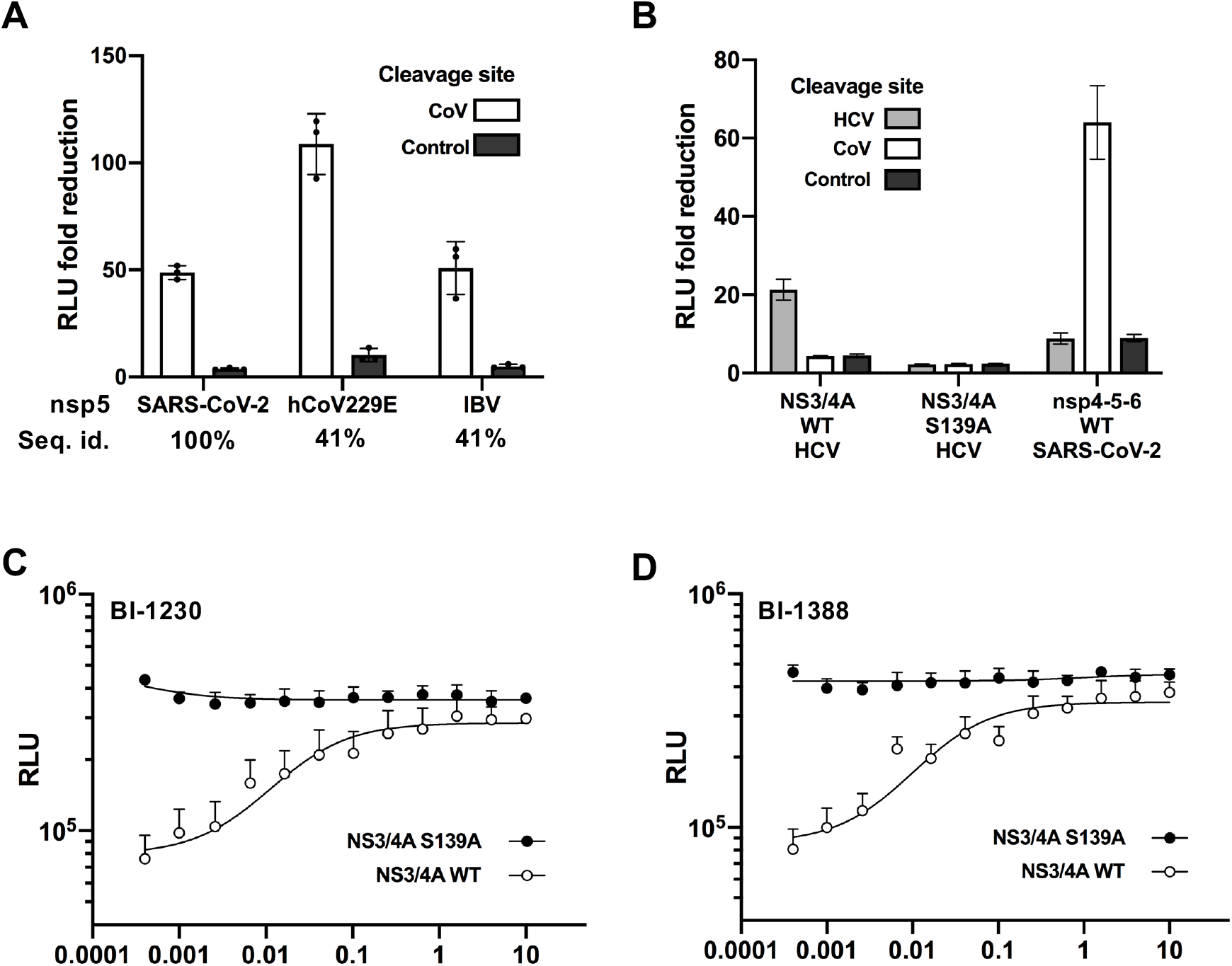
Extended use of the Rev-Nluc assay. **A**. Luciferase signal measured at 48 hpt in HEK-293T cells co-transfected with a Rev-Nluc reporter (with a CoV or control linker sequence) and nsp5 (derived from SARS-CoV-2, hCoV-229E, or IBV). The graph shows the RLU fold-reduction compared to cells transfected with an empty control plasmid (mean ± SD of three independent experiments performed in technical triplicates). The amino acid sequence identity compared to SARS-CoV-2 nsp5 is indicated below the graph. **B**. Luciferase signal measured at 48 hpt in HEK-293T cells co-transfected with a Rev-Nluc reporter (with an HCV-NS5A/5B, a CoV cleavage site, or a control linker sequence) and HCV-NS3/4A protease (WT or S139A) or SARS-CoV-2 nsp4-5-6. The graph shows the RLU fold-reduction compared to cells transfected with an empty control plasmid instead of an NS3/4A or nsp4-5-6 plasmid (mean ± SD of three independent experiments performed in technical triplicates). **C-D**. The inhibition of HCV-NS3/4A protease by BI-1230 (C) and BI-1388 (D) at concentrations from 10 to 0.0004 µM was assessed using the Rev-Nluc-based assay. The RLU are shown as the mean ± SD of two independent experiments performed in technical triplicates. Open symbols: NS3/4A WT; closed symbols: NS3/4A-S139A.

Next, we asked whether the Rev-Nluc reporter could be used to monitor the activity of a distinct class of proteases, the hepatitis C virus (HCV) NS3/4A serine protease. The Rev-Nluc-HCV reporter plasmid, in which the two Nanoluciferase subdomains are linked by the NS5A/5B cleavage site EDVVCCSMSY, was co-transfected with an expression vector for NS3/4A protease. Cells co-transfected with the NS3/4A plasmid showed a ∼20-fold reduction in luciferase signal at 48 hpt, whereas luciferase signals in cells co-transfected with the catalytically inactive S139A mutant [38] showed no change (**Figure 2B**, black bars, and **Supplemental Figure 3**). In the presence of two potent NS3/4A protease inhibitors (BI-1230 and BI-1388, Boehringer Ingelheim), a dose-dependent increase of the Nanoluciferase signal was observed in the presence of NS3/4A WT (open symbols) while the luciferase signal measured in the presence of NS3/4A S139A remained unchanged (closed symbols) (**Figure 2C-D**). The estimated IC_50_ as measured by the Rev-Nluc assay was 0.02 µM, in the same range as previously reported (https://opnme.com/).

### Implementation of an automated 384-well Rev-Nluc-assay pipeline

To allow the screening of large compound libraries with potential anti-nsp5 activity, we adapted the Rev-Nluc assay to the 384-well format in an automated pipeline as described in detail in the Methods section. In brief, HEK-293T cells are transfected in bulk, harvested at 6 hpt and distributed homogeneously in 384-well plates which contain pre-distributed compounds (**Figure 3A**). This reduces transfection efficiency variations between wells and increases time- and cost-effectiveness. In a preliminary experiment, we assessed the reproducibility of the luciferase signal across ten 384-well plates with GC376 at final concentrations of 10, 1 or 0.1 µM. The luciferase signal distribution and relative nsp5-inhibition by GC376 remained stable throughout the series, which demonstrates the applicability of the Rev-Nluc assay for high-throughput screenings (**Supplemental Figure 4**). When the same conditions were used to perform dose-response curves with CG376, Calpain Inhibitor XII, Boceprevir and the alpha-ketoamide inhibitor Mpro 13b, the relative levels of nsp5 inhibition and estimated IC_50_ values were consistent with previous reports [13, 17, 27] (**Figure 3B**). Overall, our data demonstrate that the Rev-Nluc-based is scalable to the 384-well format and can be reliably used to screen large libraries for small molecule inhibitors of SARS-CoV-2 nsp5.

**Figure 3.**
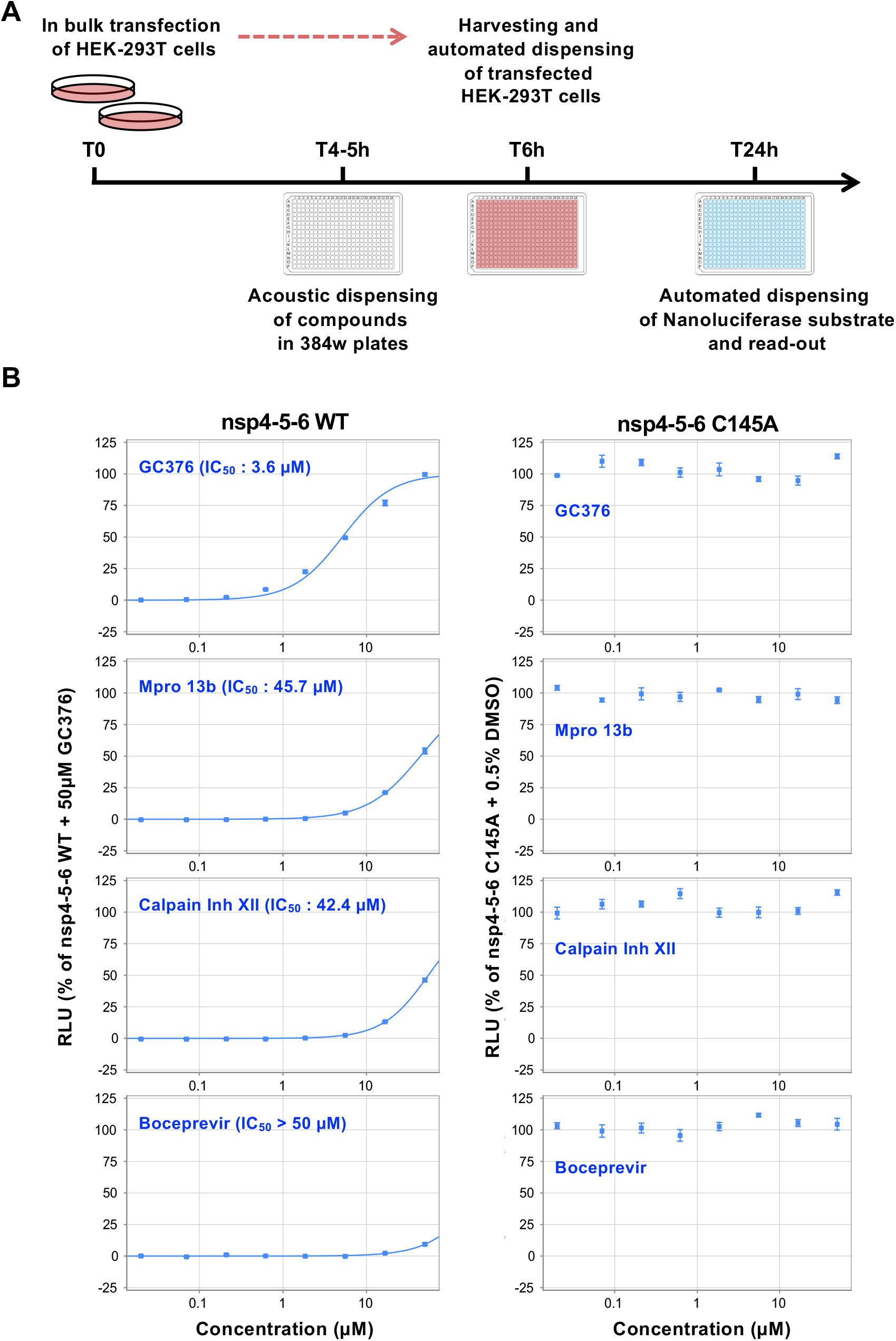
Upscaling of the Rev-Nluc-based assay to the 384-well format. **A**. Schematic representation of the automated high-throughput pipeline to perform the Rev-Nluc assay in a 384-well format. **B**. The inhibition of SARS-CoV-2 nsp5 by GC376, Mpro13b, Calpain Inhibitor XII and Boceprevir was assessed using the Rev-Nluc-based assay in a 384-well plate setting. Cells were transfected either with SARS-CoV-2 nsp4-5-6 WT (left panels) or C145A (right panels). The inhibitor concentrations correspond to 3-fold serial-dilutions from 50 to 0.02 µM. The graphs show Relative Light Units (RLU) normalized as described in the methods section (mean ± SD of technical triplicates, one representative experiment of three independent experiments is shown for GC376, Calpain Inhibitor XII and Boceprevir, two independent experiments for Mpro13b).

### *In silico*-based selection of potential anti-nsp5 compounds

We used the available structural information on ligand-bound nsp5 proteins (98 covalent and 40 non-covalent crystallographic complexes) to derive a pharmacophore (Supplemental Methods section and **Supplemental Figure 5**). Among the non-covalent complexes, 38 were bearing proper ligands to perform crossdocking experiments and were used as candidates to select for the most suitable docking protocol/schemes. The docking of an in-house library of 14,468 compounds was performed on the PDB entries 7JU7, 5R7Z and 5RH8 with the Smina software [39] and on PDB 5RH8 with the FlexX software [40], as the combination of these four docking schemes gave the best results for crossdocking on the non-covalent crystallographic complexes. Molecules which could not be docked because they had unusual atoms, had too many rotatable bonds, or could not fit for other reasons were discarded. Out of the 14,468 compounds, 13,781 could be docked at least once for one of their ∼60,000 tautomers/conformers. With up to 10 poses recorded, this led to ∼2,400,000 poses. For each docking pose, a pharmacophoric score (named PHARMscore) was calculated as described in the Methods section. Noticeably, comparison of the docking poses of FlexX and Smina on PDB 5RH8 showed better PHARMscore for FlexX. Correlations between the PHARM scores and rescoring values were significant for the selected molecules (from 0 to 58%) (“Whole selection” in the **Supplemental Table 1**). The correlation was higher for the best rescored poses than for the best PHARM poses, suggesting that the rescoring considers the molecular contacts used to calculate the PHARMscores and is more exhaustive.

**Table 1.**
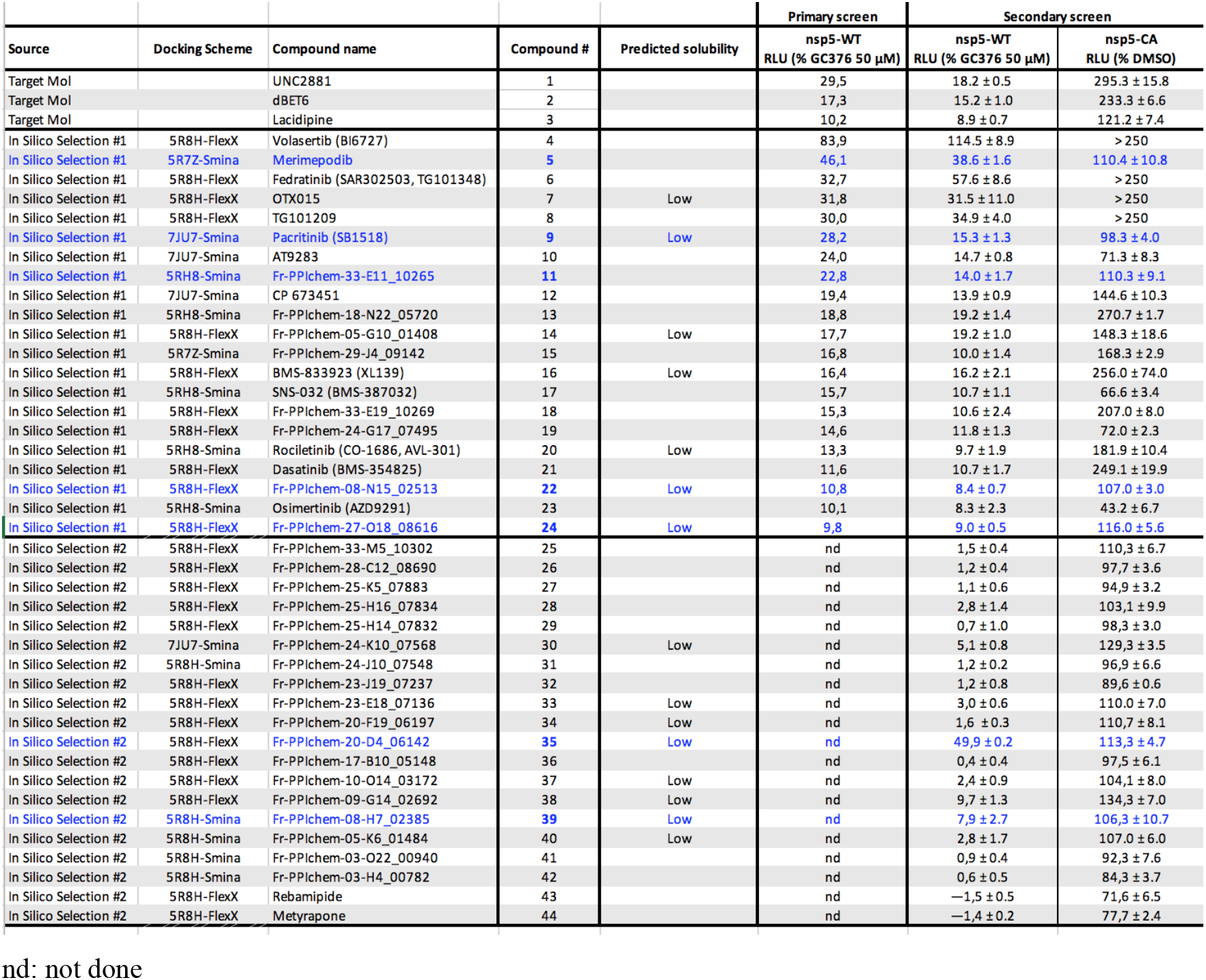
Primary and secondary screen of candidate nsp5 inhibitors.

For each molecule and each docking scheme, the pose with the top score upon docking and the pose with the top PHARMscore were selected and were used for the selection/prioritization of molecules as described in the Supplemental Methods section. This led to 393, 381, 376, 310 unique poses (or 371, 307, 319 and 371 unique molecules) for 5R7Z-Smina-HYDE, 5RH8-FlexX, 5RH8-Smina-AD4 and 7JU7-Smina-AD4, respectively, representing a total of 1460 unique poses (or 933 unique molecules) that were further analysed (**Supplemental Figure 6**). Noticeably, the following molecular interactions were observed most frequently (frequency in brackets): HBond:GLU166 (75%), Cation-π:HIS41 (46%), π-π:HIS41 (37%), HBond:GLY143 (30%), Tstack:HIS41 (27%), HBond:HIS163 (23%), SaltBridge:GLU166 (21%). In comparison, the most frequent interactions observed in the crystal complexes were HBond:GLU166 (37%), HBond:HIS163 (26%), Tstack:HIS41 (17%), HBond:GLY143 (8%). After visual inspection, 46, 121, 107 and 84 poses were selected from the 5R7Z-Smina-HYDE, 5RH8-FlexX, 5RH8-Smina-AD4 and 7JU7-Smina-AD4 docking models respectively, representing a list of 316 unique candidate molecules. An additional 23 molecules were selected because of their chemical similarity to control compounds (GC-376, Calpain inh XII, Boceprevir and masitinib), bringing to 339 the number of selected molecules (named *in silico Selection #1* below, in the **Supplemental Table 1** and **Table 1**). A second filtering of the ∼2,400,000 poses was performed based on features frequently observed both in available crystallographic complexes [9, 10, 13, 14, 17, 22, 41-43] and in the poses of the *in silico Selection #1* (namely hydrogen bonds to residues 163 and 166 and pyridine, acetamide/urea moieties) and geometric/positional similarity to crystal complexes. The 1354 filtered poses, representing 1142 unique molecules, were visually inspected to select 20 additional molecules (named *in silico Selection #2*). In total, 359 molecules (*Selection #1* and *#2*) were selected for experimental testing.

### Screening of candidate nsp5 inhibitors

We applied the 384-well Rev-Nluc-assay pipeline to test the 339 candidate nsp5 inhibitors corresponding to the *in silico Selection #1*. In parallel, we also tested a commercial (TargetMol) library of 161 small molecules with potential anti-nsp5 activity as predicted by molecular docking. In a primary screen, the compounds were distributed into 384-well plates at final concentrations of 10 µM. The luciferase signals were background subtracted and expressed as percentages of the mean signal of cells treated with the control compound GC376 (50 µM). Nsp5 inhibition of the 500 tested compounds was ranked by their potency from left to right, and the compounds showing a luciferase signal ≥10 % of the signal measured in the presence of 50 µM GC376 are represented in blue (**Figure 4A)**. Among those, 3 belong to the TargetMol library and 21 to the *in silico Selection #1* and (**Figure 4A**, light and dark blue respectively - **Table 1**, compounds #1 to #24).

**Figure 4.**
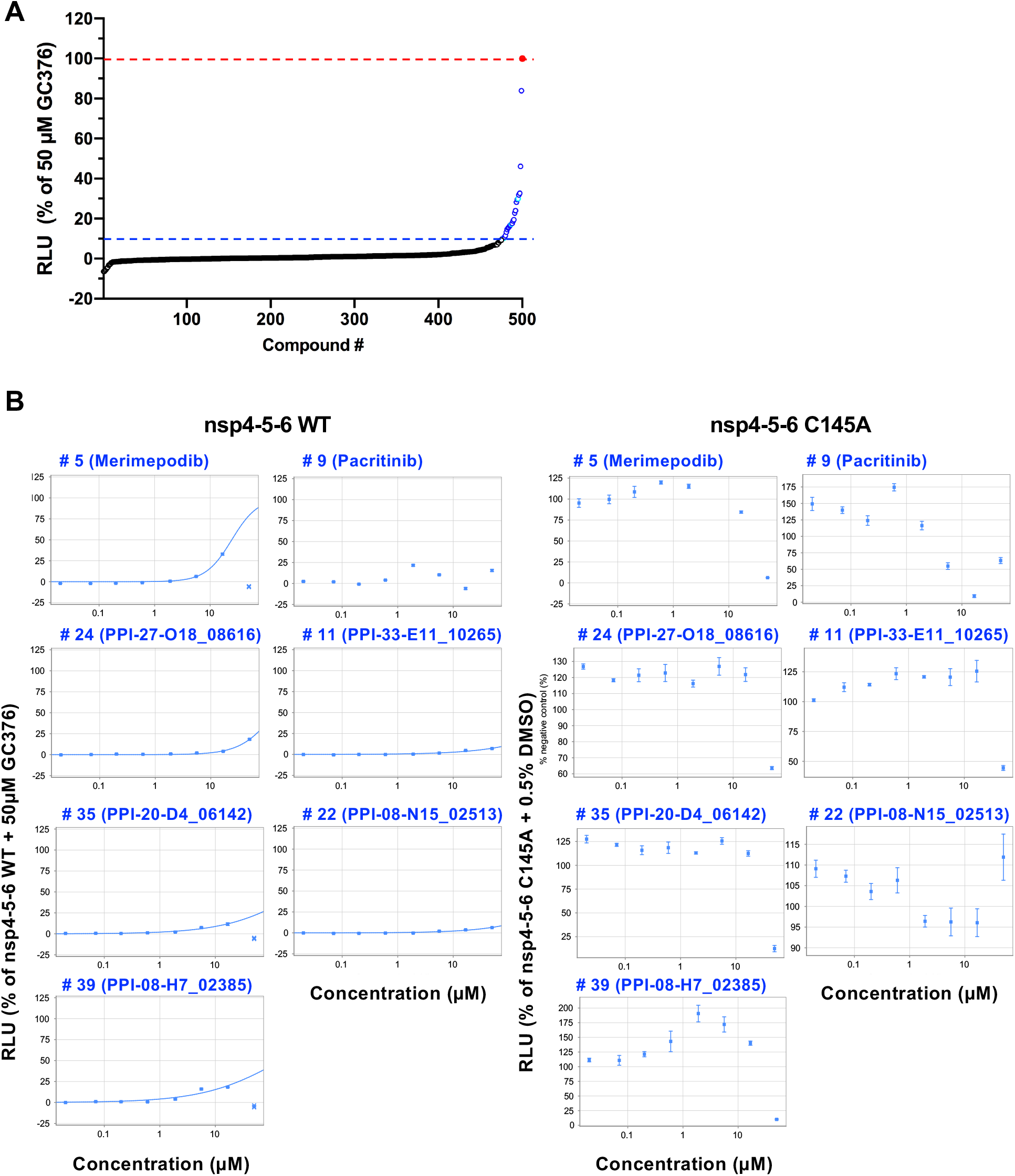
Rev-Nluc-based screening of candidate nsp5 inhibitors. **A**. Primary screen of >500 putative nsp5 inhibitors in cells transfected with SARS-CoV-2 nsp4-5-6-WT. The 339 compounds from the *in silico Selection #1* and 161 compounds from the TargetMol nsp5-targeted library were distributed into 384-well plates to achieve final concentrations of 10 µM. The graphs show Relative Light Units (RLU) normalized as described in the methods section. Compounds are ranked according to the observed increase in luciferase signal from left to right, and the primary hits (≥10 % luciferase signal compared to 50 µM GC376) are indicated in dark blue (*in silico Selection #1* compounds) or light blue (TargetMol compounds). **B**. Dose response curves on the secondary hit selection of seven compounds. Compounds #5, #9, #11, #22, #24, #35 and #39 were tested at concentrations corresponding to 3-fold serial-dilutions from 50 to 0.02 µM on cells expressing either SARS-CoV-2 nsp4-5-6 WT or C145A. The mean RLU was background subtracted (i.e. minus the signal in cells DMSO treated which were transfected with SARS-CoV-2 CoV-2-nsp4-5-6 WT). The graphs show RLU normalized as described in the methods section (mean ± SD of technical triplicates, one representative experiment of two independent experiments is shown).

To assess the specificity of nsp5 inhibition and potential cell toxicity, the 24 primary hit compounds were tested at a final concentration of 10 µM in cells transfected with either SARS-CoV-2 nsp4-5-6 WT or C145A. Compounds that led to an increased (> 120%) or decreased (< 80%) luciferase signal in SARS-CoV-2 nsp4-5-6 C145A-transfected cells were discarded. The 5 remaining compounds (# 5, 9, 11, 22 and 24, highlighted in blue in **Table 1**) induced a moderate to strong increase in luciferase signal with nsp5-WT, consistent with the primary screening data. The strongest increase (39% compared to the 50 µM GC376 control) was observed with compound #5. The twenty *in silico Selection #2* compounds were tested in the same conditions as the 24 primary hits and assessed using the same criteria, which led to the selection of two additional compounds (#35 and #39 in **Table 1**).

Dose-response curves were generated for the 7 secondary hit compounds on cells expressing either nsp5 WT or the nsp5 C145A mutant (**Figure 4B**, left and right panels respectively). The IC_50_ was estimated < 50 µM for compounds #5 (Merimepodib), #24 (Fr-PPIChem-27-O18_08616), #35 (Fr-PPIChem -20-D4_06142) and #39 (Fr-PPIChem-08-H7_02385). All four compounds became toxic at 50 µM, as indicated by a sharp decrease of the luciferase signal both on the WT and mutant nsp5. For compound #39 and to a lesser extent for compound #5, the dose-response curve on the nsp5 C145A mutant was bell-shaped with a dose-dependent increase of the luciferase signal in the 0.02 to 0.6 µM range followed by a decrease at higher concentrations, likely indicative of a non-specific effect of these compounds on the Nanoluciferase read-out combined with cell toxicity at high concentrations.

### Antiviral activity of candidate nsp5 inhibitors

The ability of compounds #5, #24, #35 and #39 (whose structure is shown in **Figure 5A**) to inhibit SARS-CoV-2 replication was evaluated in a multicycle replication assay on A549-ACE2 cells, followed by RT-qPCR quantification of the viral genomic RNA in the supernatant collected at 72 hpi (**Figure 5B**). Cytotoxicity of the compounds was assessed in parallel using an assay based on ATP quantification (**Figure 5C**). For the positive control GC376 we determined an IC_50_ of 14.7 µM and a IC_50_ to CC_50_ ratio (selectivity index or SI) > 3. Merimepodib (#5) showed an IC_50_ of 21 µM and a low SI of 1.5. Compounds #24, #35 and #39 showed a higher antiviral activity than GC376, with IC_50_ of 7.2, 12.4 and 4.7 µM and SI >7, >3 and > 10, respectively. In summary, the screening of 500 potential SARS-CoV-2 nsp5 inhibitors by the here described assay led to the identification of 2 potential lead compounds (#24, #39) with low cell toxicity and a higher antiviral activity against SARS-CoV-2 replication as compared to the reference compound GC376, which demonstrates the potential of the Rev-Nluc assay for the identification of novel SARS-CoV-2 nsp5 inhibitors.

**Figure 5.**
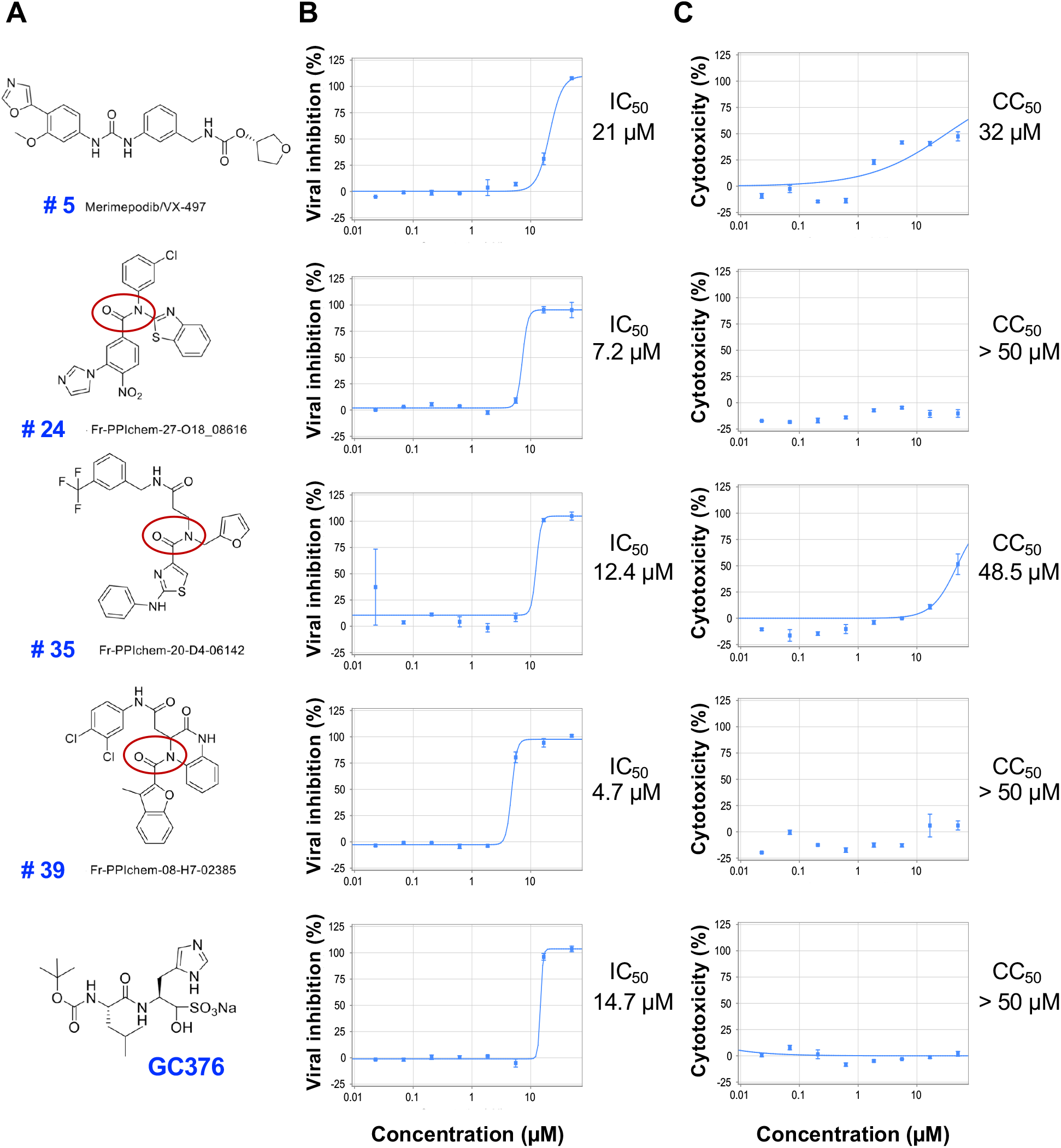
Antiviral activity and cytotoxicity of candidate nsp5 inhibitors. **A**. The structure of the four secondary hit compounds (#5, #24, #35 and #39) is shown and their tertiary amide component is circled. GC376 was used as a control. **B**. The ability of compounds #5, #24, #35 and #39 to inhibit SARS-CoV-2 replication was evaluated at concentrations corresponding to 3-fold serial-dilutions from 50 to 0.02 µM in a multicycle replication assay on A549-ACE2 cells, followed by RT-qPCR quantification of the viral genomic RNA in the supernatant collected at 72 hpi. The percentages of viral inhibition was determined as described in the Methods section. (one representative experiment of two independent experiments is shown). **C**. In parallel the indicated compounds were tested on A549-ACE2 cells for cell cytotoxicity by the CellTiter-Glo assay. In B and C, the data are expressed as the mean ± SD of technical triplicates (one representative experiment of two independent experiments is shown).

## Discussion

We have developed and validated a sensitive high-throughput cell-based assay to monitor SARS-CoV-2 nsp5 proteolytic activity. We screened a library of molecules for SARS-CoV-2 nsp5 inhibitors, and identified 4 novel lead compounds. This was achieved by engineering a Reverse-Nanoluciferase (Rev-Nluc) reporter in which two Nanoluciferase domains are permuted and linked together by an nsp5 cleavage site. Co-expression with the SARS-CoV-2 nsp4-5-6 polyprotein results in cleavage of the reporter and thereby a reduced luciferase activity. The addition of a specific nsp5 inhibitor results in a dose-dependent increase in luciferase activity. A ∼30-fold increase in luciferase activity is observed in the presence of 50 µM GC376, similar to the ∼30-fold increase observed when a control inactive nsp4-5-6 polyprotein is co-expressed with the reporter instead of the wild-type nsp4-5-6. The strengths of the Rev-Nluc-based assay lie in the following: i) it is a gain-of-signal assay and therefore excludes compounds that are cytotoxic or interfere negatively with the Nanoluciferase read-out; ii) it is rapid and convenient due to the glow-type signal of the Nanoluciferase; iii) it is amenable to miniaturisation due to the very bright luminescence signal produced by the Nanoluciferase; and iv) as a consequence of its rapidity and miniaturisation it can be run in parallel on a wild-type and a catalytically inactive nsp5, which allows for the identification of compounds that enhance the luciferase activity in both conditions and to discard them as false-positives. Unlike previously described SARS-CoV-2 nsp5 reporter assays, we provide evidence that our Rev-Nluc-based assay is scalable for high-throughput screens in a 384-well format.

The first cell-based assays for SARS-CoV-2 nsp5 to be described were loss-of-signal assays based on a Flip-GFP [26] or a Flip-Firefly [31] reporter. They are less scalable for high-throughput application than the Rev-Nluc-based assay because of the low intensity and narrow dynamic range of fluorescence signals, or the flash-type signal of the Firefly luciferase. To date, two gain-of-signal assays were described. The first assay relies on a chimeric protein containing the eGFP reporter and the full nsp5 amino acid sequence flanked by cognate self-cleavage sites [29]. The mechanism underlying the fluorescent read-out is unclear and the scalability of the assay is limited. The second assay uses crystal violet staining as a read-out to monitor cytotoxicity upon nsp5 overexpression and is therefore not specific for nsp5 protease inhibition, as reduced cytotoxicity may result from off-target effects of the compounds [30]. More recently Rawson et al. described a gain-of-signal assay [27], which differs from ours in that it is based on the NanoBiT system [44]. The two Nanoluciferase-derived domains which are linked together by an nsp5 cleavage site are more unbalanced in size (156 and 13 amino acids, compared to 105 and 61 for Rev-Nluc) and together they show 15 amino acid substitutions compared to the Rev-Nluc sequence. The ability of the NanoBiT-based assay to identify nsp5 inhibitors was demonstrated in a 96-well format, however its performance in terms of stability and scalability for a high-throughput screens was not investigated [27].

We scaled-up the Rev-Nluc assay to an automated 384-well format, and used these conditions to assess the inhibition of SARS-CoV-2 nsp5 by a set of reference compounds: GC376, boceprevir and calpain inhibitor XII, which have been shown to inhibit nsp5 and to impair viral replication (e.g. [17, 22, 23, 36, 43], and the alpha-ketoamide inhibitor 13b [13]. Our findings and IC_50_ estimations (**Figure 3B**) were consistent with previous reports [13, 17, 27], therefore validating our assay.

We screened a total of 520 molecules with potential anti-nsp5 activity as predicted by molecular docking: 359 molecules from the *in silico* screening approach and 161 molecules from a commercial targeted library. A primary screen, performed at a single concentration on the wild-type nsp5 protein only, identified 24 hits. The subsequent secondary screen, performed at a single concentration on the wild-type and catalytically inactive nsp5 proteins in parallel, led us to discard most of the primary hits as being toxic or false-positives and to select only 7 molecules, all coming from the *in silico* screening approach. These results highlight the fact that being able to manage parallel screening on the wild-type and CA nsp5 proteins provides a major added value to our assay in terms of selectivity. Moreover, the fairly high hit rate of 7 hits out of 359 molecules demonstrates the effectiveness of the *in silico* screening approach, characterized by i) a careful selection of the available nsp5 structures to be used for docking, ii) a special attention paid to the *in silico* preparation of ligands for docking, iii) a comparison of several docking protocols, iv) the complementary use of a pharmacophoric score based on available structural information, and v) a precise analysis of the poses.

Upon dose-response analysis of the 7 selected hits, 4 molecules showed an estimated IC_50_ below 50 µM in the Rev-Nluc assay and were therefore assessed in a multicycle SARS-CoV-2 replication- and a cytotoxicity-assay. These molecules include merimepodib/VX-497, an inhibitor of the human inosine monophosphate dehydrogenase initially developed as an immunosuppressor [45]. Merimepodib was later found to have antiviral activity against several DNA and RNA viruses including HCV, Zika, Ebola and FMDV viruses [46-49]. In the early phase of the COVID-19 pandemic, oral merimepodib was investigated in combination with intravenous remdesivir administration in a phase 2 clinical trial as a potential treatment for severe COVID-19. The trial sponsor announced in October 2020 that the trial was discontinued because it was unlikely that it would meet its primary safety endpoints [50]. Indeed, we found an EC_50_ of 21 µM with a very low selectivity index of 1.5 for merimepodib on A549-ACE2 cells. More interestingly however, our data suggest a particular and unexpected mode of action of this compound against the SARS-CoV-2 virus. Merimepodib is thought to exert its broad antiviral activity by reducing the pool of intracellular guanine nucleotides and thereby impairing RNA and DNA synthesis. Our findings indicate that it specifically inhibits SARS-CoV-2 nsp5 with an IC_50_ close to 16 µM. The question whether merimepodib can inhibit other viral cysteine-dependent proteases and to what extent this anti-protease activity contributes to its antiviral activity deserves to be further explored.

We also identified three compounds (#24, #35 and #39 in **Table 1** and **Figure 5**) which inhibit SARS-CoV-2 replication on A549-ACE2 cells more efficiently than GC376, with IC_50_ values in the 4-12 µM range and selectivity indexes between 4 and 10. These three molecules are components of the Fr-PPI-Chem library of putative protein-protein interaction inhibitors [51]. Interestingly they share a common tertiary amide component (circled on the structures shown in **Figure 5A**) as well as in series of amides reported to inhibit the SARS-CoV nsp5 protease [42, 52, 53]. Given the chemical structure of the of compounds #24, #35 and #39 and their predicted pKa and clogP values (in the 1.6-3.9 and 2.28-6.57 range, respectively), addition of a proton leading to cationic amphiphilic molecules and, as a consequence, inducing a phospholipidosis that can be confused with an antiviral effect [54], is unlikely. Interestingly, the number of hits obtained from the *in silico* screening was comparatively high, retrospectively supporting the cautious choice of virtual screening conditions made. Further experiments, including X-ray crystallography and biochemistry, will be needed to investigate the functional importance of this tertiary amide component, and to further assess the potential of compounds #24, #35, #39, or merimepodib as starting points for the design of more effective nsp5 inhibitors.

## Acknowledgments

We thank Nicholas Heaton (Duke University), Rolf Hilgenfeld (University of Luebeck), Olivier Schwartz, Sylvie van der Werf and Olivier Sperandio (all from Institut Pasteur) for providing biological material and advice. We thank Yves Janin (Institut Pasteur) for providing reagents and for insightful suggestions on the manuscript, and Mallory Perrin-Wolff (Institut Pasteur) for her continuous support.

## Figure legends

**Supplemental Table 1.** Average rescoring values and PHARM score and their correlations for poses selected by best rescoring values (best-score) and best-PHARM score (best-PHARM) (see Supplemental Methods section). The statistics are calculated on Selection #1, Selection #2, and on the merge of both selections (“Whole selection”).

| Dock\correl | Selection #1 |  |  |  |  |  |
| --- | --- | --- | --- | --- | --- | --- |
|  | best-score |  |  | best-PHARM |  |  |
|  | Av.Score | Av.Pharm | Correl | Av.Score | Av.Pharm | Correl |
| 5r7z | -22.69 | 20.11 | -0.23 | -3.61 | 31.25 | -0.10 |
| 5rh8-FlexX | -32.17 | 38.68 | -0.59 | -28.48 | 48.65 | -0.45 |
| 5rh8 | -8.53 | 29.42 | -0.35 | -7.59 | 35.63 | -0.32 |
| 7ju7 | -8.12 | 26.95 | -0.24 | -6.82 | 35.40 | -0.32 |
|  | Selection #2 |  |  |  |  |  |
|  | best-score |  |  | best-PHARM |  |  |
|  | Av.Score | Av.Pharm | Correl | Av.Score | Av.Pharm | Correl |
| 5r7z | -19.48 | 16.97 | -0.12 | 1.41 | 26.12 | 0.26 |
| 5rh8-FlexX | -31.03 | 35.24 | -0.42 | -28.29 | 43.48 | -0.35 |
| 5rh8 | -7.99 | 26.03 | -0.81 | -7.16 | 33.05 | -0.48 |
| 7ju7 | -7.43 | 27.95 | -0.17 | -6.57 | 32.56 | -0.31 |
|  | Whole Selection |  |  |  |  |  |
|  | best-score |  |  | best-PHARM |  |  |
|  | Av.Score | Av.Pharm | Correl | Av.Score | Av.Pharm | Correl |
| 5r7z | -19.43 | 17.30 | -0.21 | -4.54 | 25.87 | 0.07 |
| 5rh8-FlexX | -26.88 | 30.39 | -0.58 | -24.45 | 38.89 | -0.53 |
| 5rh8 | -5.65 | 23.58 | -0.43 | -4.81 | 29.25 | -0.38 |
| 7ju7 | -7.42 | 22.27 | -0.34 | -6.34 | 29.57 | -0.36 |

**Supplemental Table 2.**
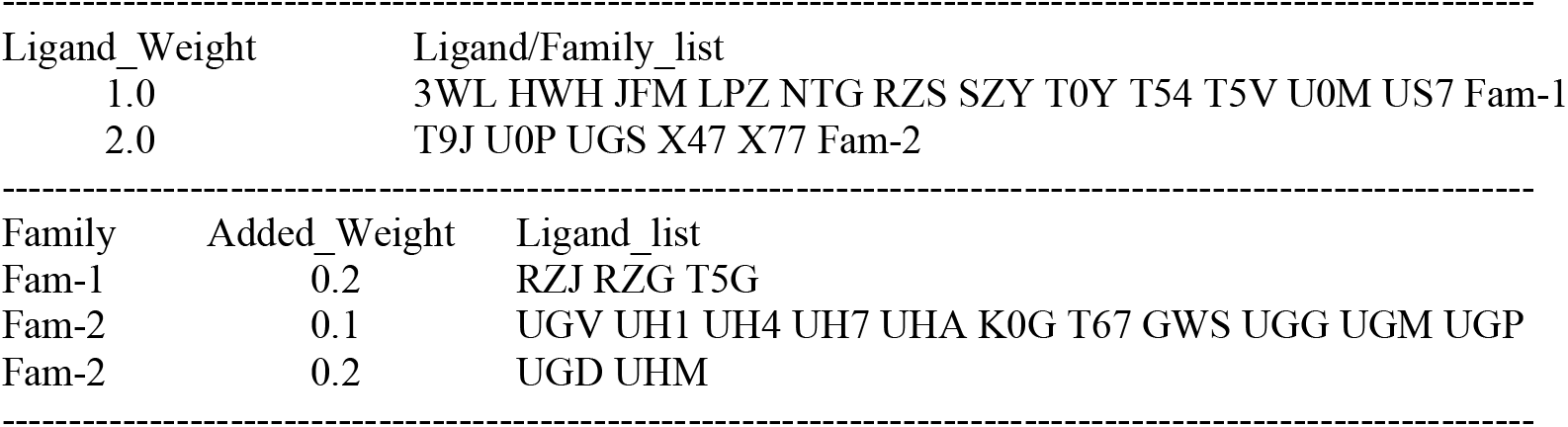
Weight for successfully docked ligands. If more than one ligand from a family of similar ligands (Fam-1 or Fam-2) appears, the weight of the family is counted once, and the additional weight of each ligand is summed.

## Legends of Supplemental Figures

**Supplemental Figure 1:**
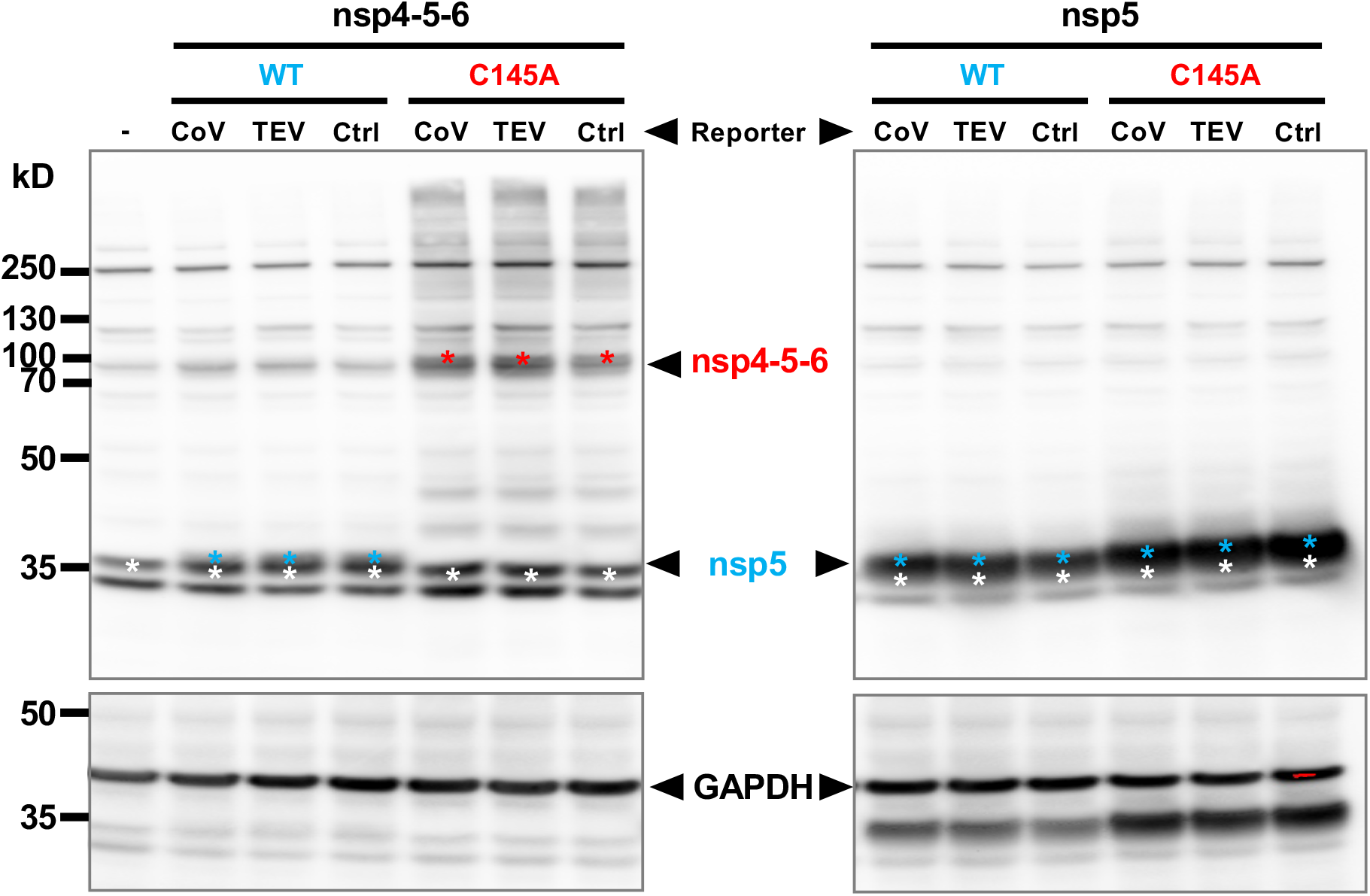
Western blot analysis of nsp5 and nsp4-5-6 expression. Expression of SARS-CoV-2 nsp4-5-6 and nsp5 in the experiment as performed in Figure 1B was assessed by western blot. HEK-293T cells were transfected with a Rev-Nluc reporter (with a CoV or TEV cleavage site, or a control linker sequence) and SARS-CoV-2 nsp4-5-6 WT or C145A. Total cell lysates were prepared at 48 hpt in Laemmli buffer and were analyzed by western blot using a polyclonal antiserum directed against SARS-CoV-2 nsp5 (upper panels) or an anti-GAPDH antibody (lower panels). Molecular weight markers (kDa) are indicated on the left. The stars point to the nsp4-5-6-specific bands (pink), nsp5-specific band (blue) and a non-specific band (white).

**Supplemental Figure 2:**
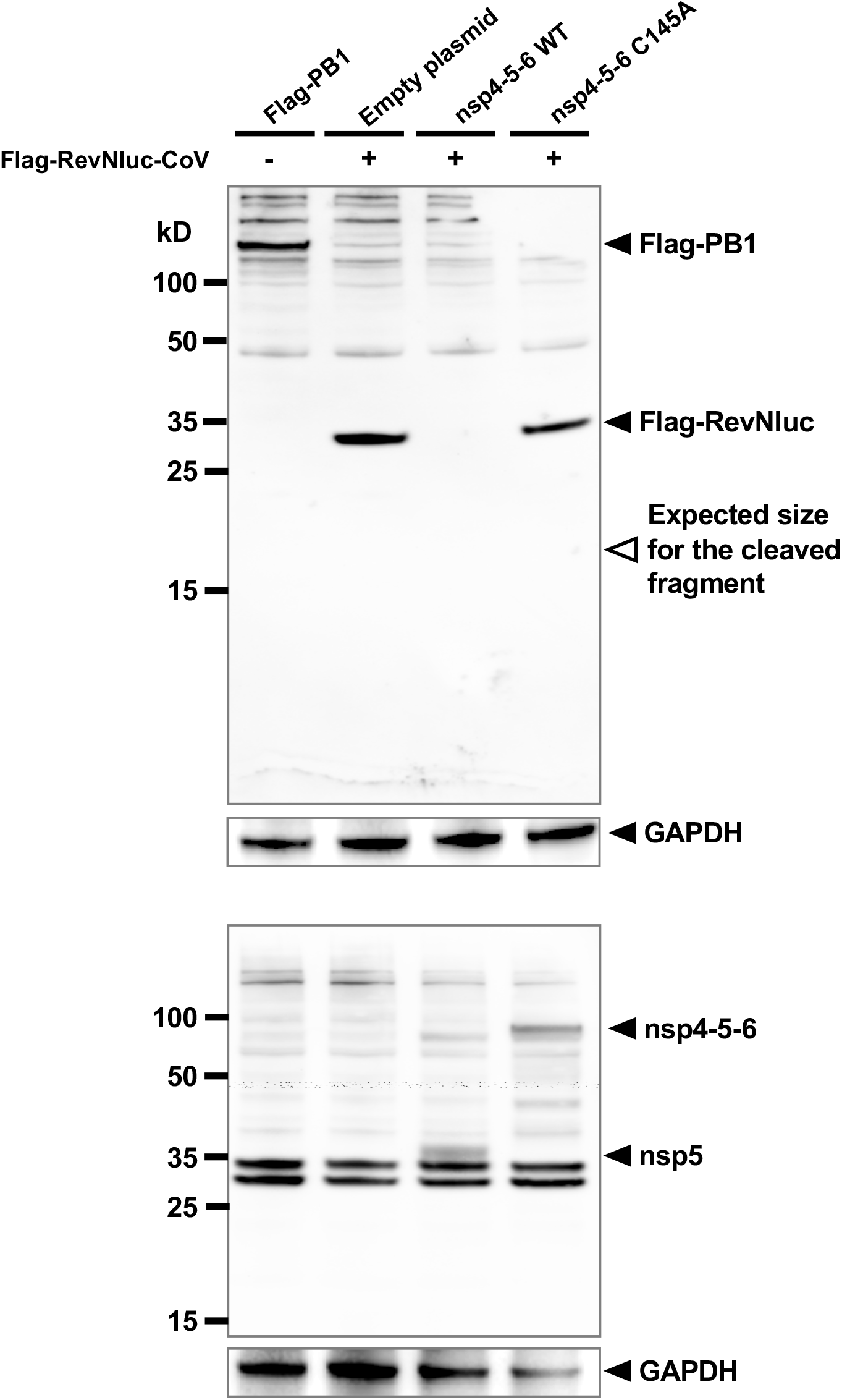
Western blot analysis of nsp5-mediated cleavage of the Flag-Rev-Nluc-CoV reporter. HEK-293T cells were transfected with the indicated plasmids. Total cell lysates were prepared at 48 hpt in Laemmli buffer and analyzed by western blot using an antibody directed against the FLAG-tag (large upper panel), SARS-CoV-2 nsp5 (large lower panel) or GAPDH (lower panels). Molecular weight markers (kDa) are indicated on the left. A Flag-PB1 (influenza Polymerase Basic 1 protein) plasmid was used as a positive control. The open arrowhead points to the expected migration of the FLAG-tagged cleavage product of Rev-Nluc-CoV.

**Supplemental Figure 3:**
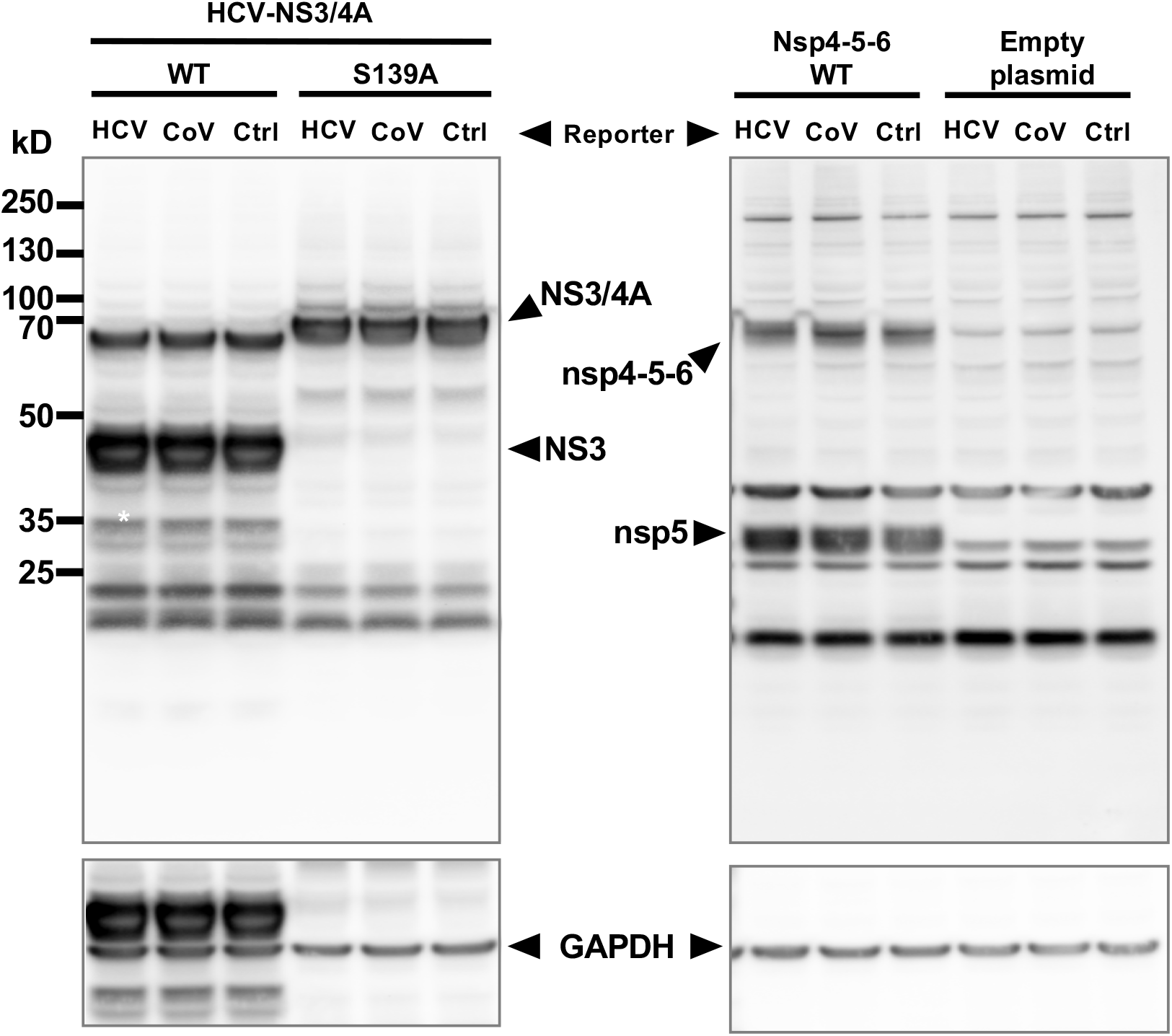
Western blot analysis of NS3/4A and SARS-CoV-2 nsp4-5-6 expression. Expression of HCV NS3/4A and SARS-CoV-2 nsp4-5-6 as performed in Figure 2B was assessed by western blot. HEK-293T cells were transfected with the indicated combinations of Rev-Nluc reporter (with a Cov or HCV cleavage site, or a control linker sequence) and HCV-NS3/4A or SARS-CoV-2 nsp4-5-6. Total cell lysates were prepared at 48 hpt in Laemmli buffer and analyzed by western blot using an antibody directed against HCV-NS3/4A or SARS-CoV-2 nsp5 (upper panels) or GAPDH (lower panels). Molecular weight markers (kDa) are indicated on the left.

**Supplemental Figure 4.**
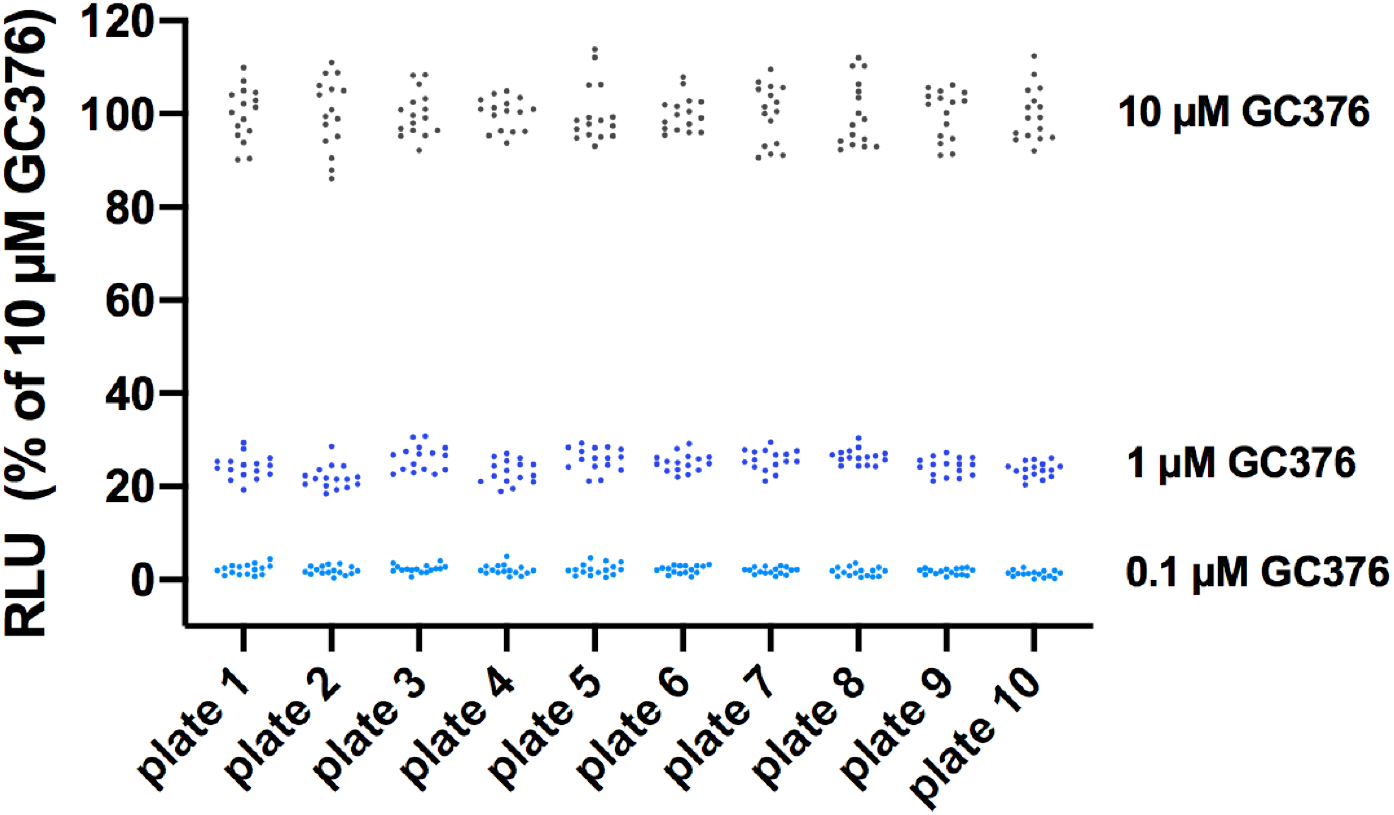
Robustness of the Rev-Nluc-based assay in the 384-well format. The Rev-Nluc assay was performed as described in the methods section throughout a series of ten 384-well plates. SARS-CoV2-nsp5 inhibition was assessed in the presence of GC376 at final concentrations of 10, 1 or 0.1 µM (16 wells for each condition in each plate). As described in Figure 3A, HEK293T cells co-transfected with Rev-Nluc-CoV and SARS-CoV-2 nsp4-5-6 WT were seeded on top of pre-distributed GC376 inhibitor. Each dot represents the luciferase signal measured in a well with 10 (black), 1 (dark blue), or 0.1 (light blue) µM GC376. The Relative Light Units (RLU) are background-subtracted (i.e. minus the signal in cells treated with DMSO) and expressed as percentages of the mean signal measured in 16 wells treated with 10 µM GC376.

**Supplemental Figure 5:**
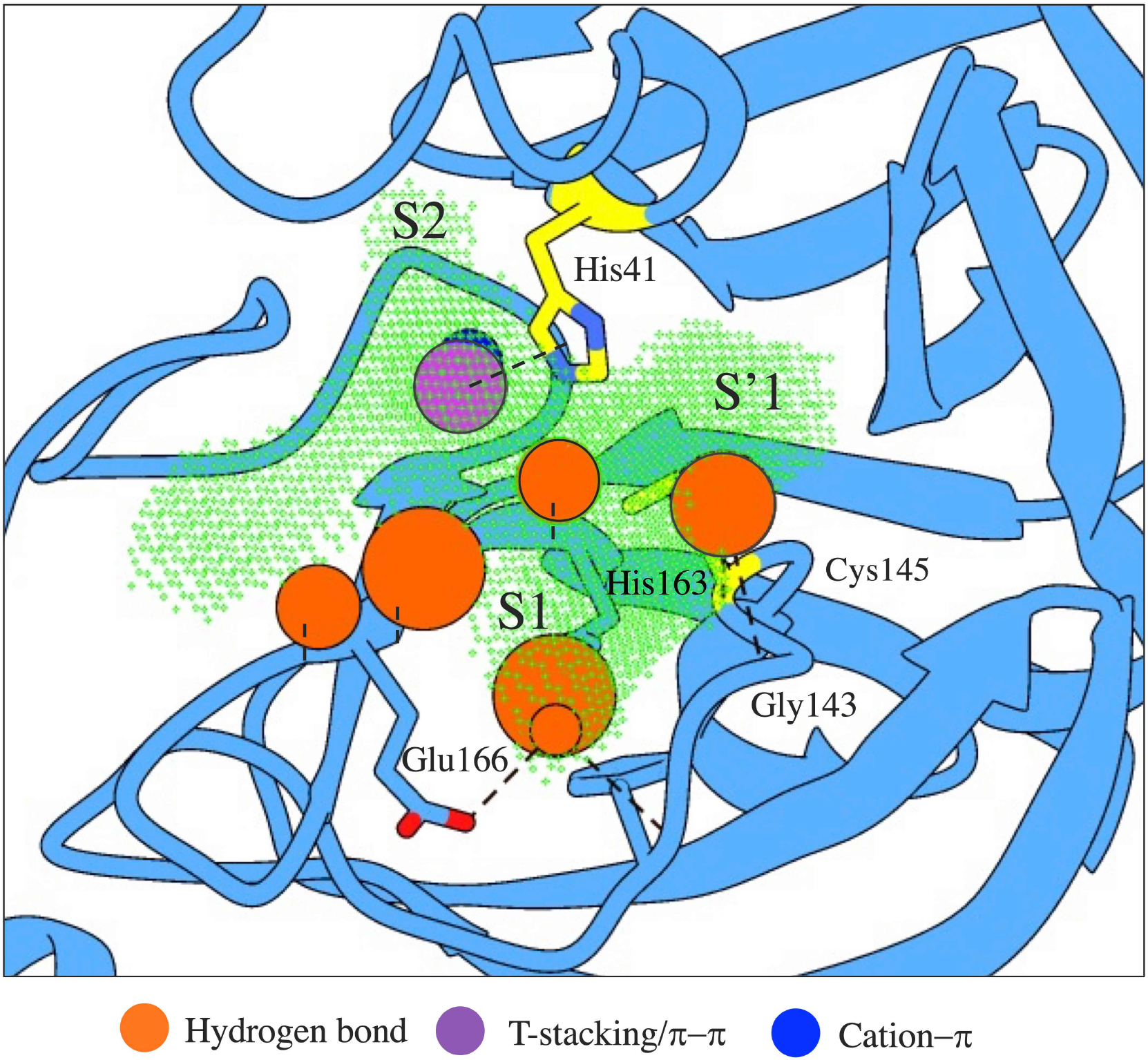
Schematic view illustrating the Pharmacophoric model. The structure of the catalytic active site of nsp5 is displayed as blue ribbons (PDB?). Carbons of the catalytic residues His41 and Cys145 are represented in yellow. Non-covalent interactions between residues (hydrogen bonds, T-stacking/π-π and cation-π interactions) are indicated by colored disks, the surface of which is proportional to the weight of the interaction. Residues that make contacts are labeled. The void volume of the catalytic groove as calculated by *mkgridXf* [55] is indicated by the dotted surface and the three corresponding sub-pockets are labelled S1, S2 and S’1.

**Supplemental Figure 6.**
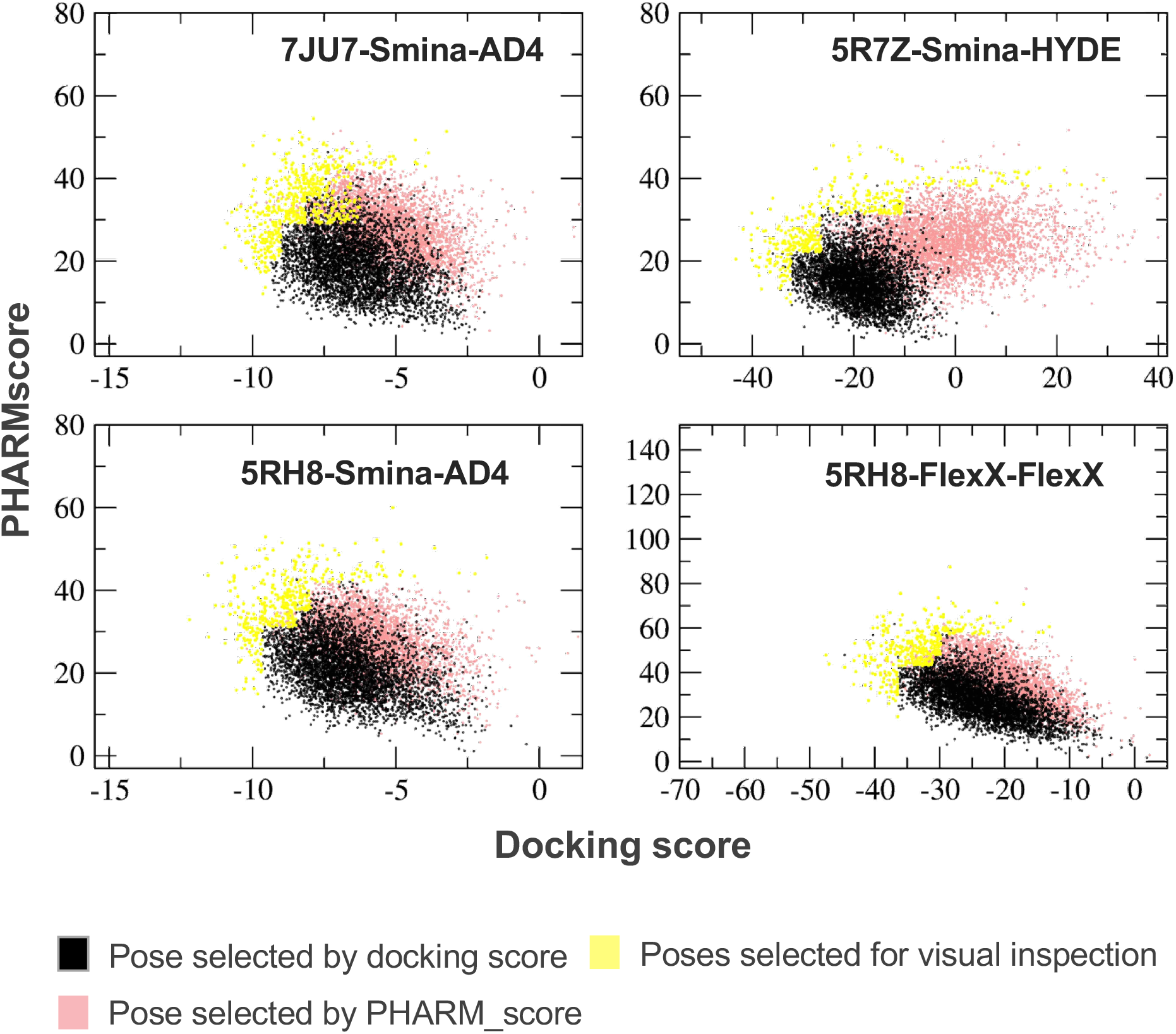
Selection of compounds based on docking pose rescoring and the PHARMscore. Top-docking score poses (black dots) and top-PHARMscore poses (salmon dots) are shown for each of the screened compounds with predicted medium/high solubility (logS and SFI calculations) and compliant with the Veber rule in each of the 4 screening protocols (denoted PDB-Sampling engine-Scoring function). The poses selected for visual inspection are highlighted in yellow.

**Supplemental Figure 7:**
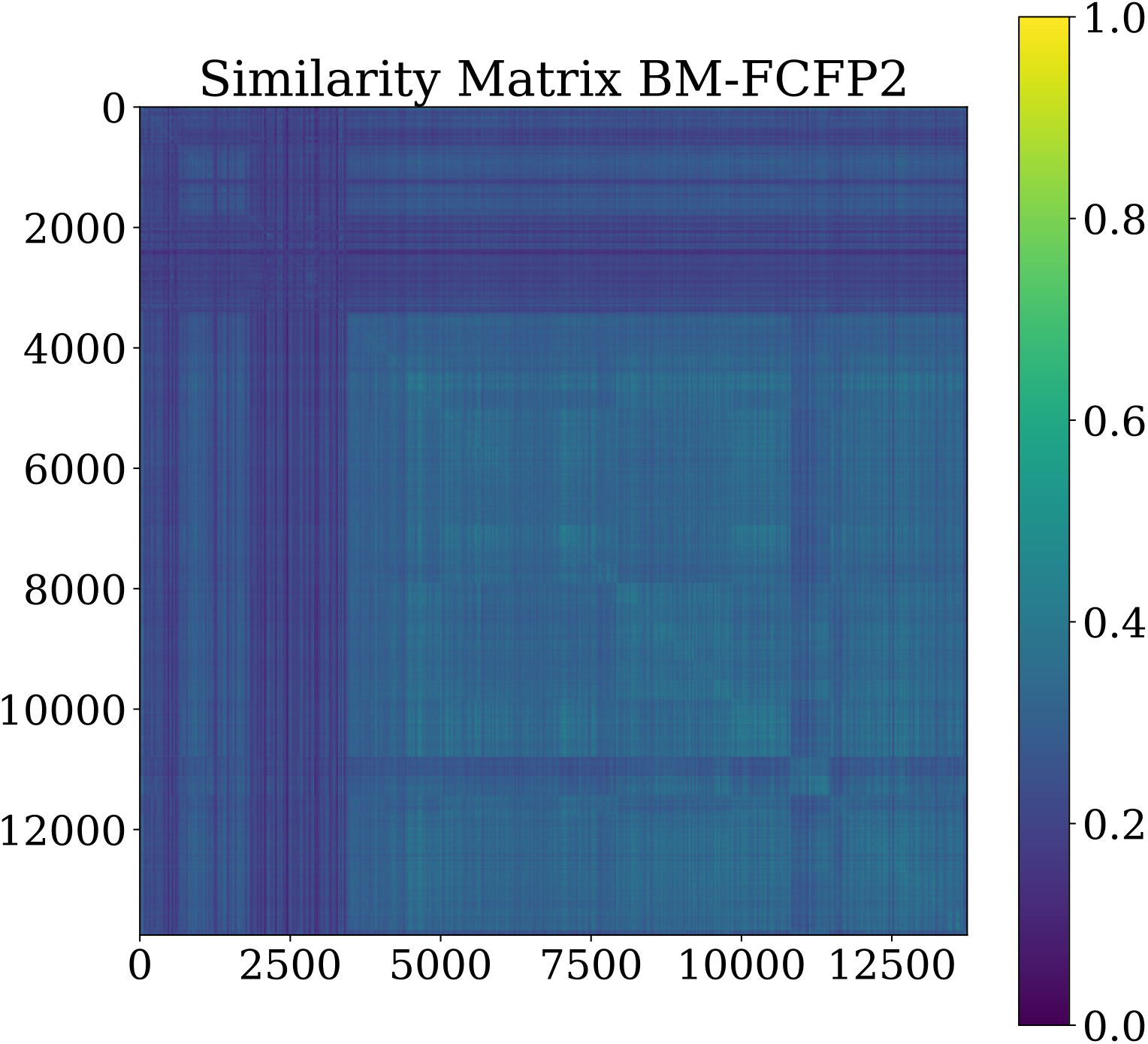
Similarity matrix for compounds of the Institut Pasteur in-house chemical library. BM-FCFP2 was used to calculate pairwise similarity among the 14,468 compounds that were included in the *in silico* screen. Similarity scores range from 1 (100% similarity, dark blue) to 0 (no similarity, yellow). The similarity matrix reveals predominantly low similarity scores, reflecting the high chemical diversity. Magnification is needed to visualize the 100% similarity scores along the diagonal of the matrix.

## Supplemental Material and Methods

### Antibodies and immunoblots

Protein extracts were prepared in Laemmli buffer. Immunoblot membranes were incubated with primary antibodies directed against nsp5 [31], NS3/4A (ab21124, Abcam), FLAG-tag (F3165, Sigma) or GAPDH (MA5-15738, Invitrogen), and revealed with HRP-linked secondary antibodies (Jackson ImmunoResearch) using ECL2 substrate (Pierce). The chemiluminescence signals were acquired with the ChemiDoc imaging system and the images were analyzed with ImageLab (Bio-Rad).

### Virtual screening for anti-nsp5 compounds

A sdf format file containing the “2D” molecule information was exported from the Collaborative Drug Discovery program (www.collaborativedrug.com) used to manage the *in-house* Pasteur chemical library. This library includes the Fr-PPI-Chem library of 10,314 putative protein-protein interaction (PPI) inhibitors [51]. The remaining 4,154 compounds (including FDA approved drugs, natural compounds, anti-cancer and anti-viral candidates) were purchased from Enzo Life Sciences, Sigma, Selleckchem, TargetMol, Greenpharma, MedChemExpress, Cayman Chemicals, Tocris, Clinisciences, or Specs. The chemical diversity of the library is illustrated in the **Supplemental Figure 7**. A “3D” sdf version was prepared for docking using an in-house automated pipeline performing: Chemical standardization and curation (ChEMBL protocols [56]); Removal of chemical duplicates (Open Babel [57]); Selection of dominant tautomers (pH 7.4, ChemAxon Calculator Plugins version 19.0.9, https://chemaxon.com); Selection of species populating more than 25% or, if none, species that are present at more than 75% of the most frequent one, or at least the three most frequent ones; Generation of stereoisomers and low energy ring conformers generation (CORINA classic [58] version 3.4, https://www.mn-am.com); Slight minimization (RDKit version 2018.03.4, https://www.rdkit.org); Annotation and flagging for solubility (ChemAxon Calculator Plugins), and for toxicophores, “Pan Assay Interference Compounds” (PAINS) [59], “Solubility Forecast Index” [60], “Quantitative Estimate of Drug-likeness” (QED) [61], and the Veber rule ([62], implemented in an in-house script using RDKit). Molecules were kept in the sdf format for docking with FlexX [40] and converted into pdbqt files using MGLTools scripts [63]for docking with Smina [39].

We selected three structures (7JU7 [64], 5R7Z [65] and 5RH8) from the Protein Data Bank (PDB, [66]) to perform the docking of these libraries. We used Smina (version Dec 2020 based on Autodock Vina 1.1.2, exhaustiveness set to 12, 10 best score poses saved) to perform docking targeting all three PDB structures. For 7JU7 and 5RH8, a 12.75 × 22.5 × 16.5 Å box centered at x = 9.02, y = -0.4, z = 25 (wrt PDB 5R7Z) was used and rescoring was made with Autodock4 ([67], AD4, version 4.2.5). For PDB 5R7Z, a box 18 × 21 × 18 Å, centered at x = 8.1, y = 0.6, z = 24.3 (wrt PDB 5R7Z) was used with rescoring with HYDE [68, 69]. FlexX was also used to perform docking with 5RH8 with the same binding site definition as for Smina (version 2.3.2, automatic selection of the base fragments, 1000 steps of optimization, 2 Å of RMSD for clustering; 10 best-score poses saved). This combination was the best of all possible combinations of four docking schemes using Smina and/or FlexX and non-covalent crystallographic complexes available in the PDB at the time to accurately perform crossdocking (see Supplemental Methods section). The nsp5 structures were prepared for docking by analysing optimal protonation states and orientation of rotatable side chains, removing all co-crystallized water and ions, and were converted into ready-to-dock formats using Open Babel and MGLTools scripts.

Beyond the rescoring values, all docking poses were further characterized by a pharmacophoric score (herein called PHARMscore), derived from available co-crystal structures of nsp5 with covalent and non-covalent ligands. It was calculated as the sum of the ligand-receptor molecular interactions detected by the BINANA algorithm [70], sorted in two categories, non-specific (2.5 Å contacts, 4 Å contacts, and hydrophobic contacts) and specific (hydrogen bonds, salt bridges, cation-π, π-π stacking and T-stacking interactions), normalized for the average contact count per interaction type, and for the specific set, weighted according to their observed frequency in the crystallographic complexes available in the PDB at the time (a schematic view of the pharmacophore is provided in the **Supplemental Figure 6)**. The top-rescored poses and the top-PHARMscore poses for each of the screened compounds (∼14 000) and for each of the 4 screening protocols were retained. Selections of poses was made by union of 4 channels per screening campaign: 1-the 150 best PHARMscore from the top 600 molecules from the best rescored list; 2-the 150 best rescored from the top 600 molecules from best PHARMscore list; 3-the top 100 molecules form the best rescored list; 4-the top 100 molecules from the best PHARMscore list. Key interactions (hydrogen bond with His163 and Glu166) and chemical/positional similarity to complexed ligands were used to prioritize compounds compliant with Veber rules and not having a low calculated solubility nor a solubility forecast index higher than 5. Some molecules that did not comply with these solubility/availability rules were still selected. All poses selected at this point were visually inspected to refine selection for experimental testing.

MolGpka, a graph-convolutional based method for pKa calculation (https://xundrug.cn/molgpka) [71], was used to calculate the pKa of hit molecules, and the Molinspiration software (https://www.molinspiration.com/) was used to predict their octanol/water partition coefficient (cLogP).

### Covalent crystallographic complexes used for analyses

5REJ, 5REK, 5REL, 5REM, 5REN, 5REO, 5REP, 5RER, 5RES, 5RET, 5REU, 5REV, 5REW, 5REX, 5REY, 5RFF, 5RFG, 5RFH, 5RFI, 5RFJ, 5RFK, 5RFL, 5RFM, 5RFN, 5RFO, 5RFP, 5RFQ, 5RFR, 5RFS, 5RFT, 5RFU, 5RFV, 5RFW, 5RFX, 5RFY, 5RFZ, 5RG0, 5RG2, 5RG3, 5RGL, 5RGM, 5RGN, 5RGO, 5RGP, 5RGT, 5RH5, 5RH6, 5RH7, 5RH9, 5RHA, 5RHB, 5RHC, 5RHE, 5RHF, 6LU7, 6LZE, 6M0K, 6W2A,, 6WNP 6WTJ, 6WTK, 6WTT, 6XA4, 6XB2, 6XBG, 6XBH, 6XBI, 6XCH, 6XFN, 6XHL, 6XHM, 6XHN, 6XHO, 6XMK, 6XQS, 6XQT, 6XQU, 6XR3, 6Y2F, 6Y2G, 6YNQ, 6YT8, 6YZ6, 6ZRT, 6ZRU, 7BQY, 7BRP, 7BRR, 7BUY, 7C6S, 7C6U, 7C7P, 7C8B, 7C8R, 7C8T, 7C8U, 7CBT, 7COM, 7JYC

### Non-covalent crystallographic complexes used for analyses

5R7Y, 5R7Z, 5R80, 5R81, 5R82, 5R83, 5R84, 5RE4, 5RE9, 5REB, 5REH, 5REZ, 5RF1, 5RF2, 5RF3, 5RF6, 5RF7, 5RFE, 5RG1, 5RGH, 5RGI, 5RGK, 5RGU, 5RGV, 5RGW, 5RGX, 5RGY, 5RGZ, 5RH0, 5RH1, 5RH2, 5RH3, 5RH8, 5RHD, 6M2N, 6W63, 6W79, 6WCO, 7JU7. 5RF2 was only used to assess contacts for the Pharmacophore.

### Structure selection for virtual screening

Analysis of the available non-covalent complex structures revealed very close conformations (making a first cluster: 5R7Y, 5R7Z, 5R80, 5R81, 5R82, 5R83, 5R84, 5RE4, 5RE9, 5REB, 5REH, 5REZ, 5RF1, 5RF3, 5RF6, 5RF7, 5RFE, 5RG1, 5RGH, 5RGI, 5RGK, 5RGU, 5RGV, 5RGW, 5RGX, 5RGY, 5RGZ, 5RH0, 5RH1, 5RH2, 5RH3, 5RH8, 5RHD, within 0.5 Å RMS on common atoms) and a few outliers (6M2N, 6W63, 6W79, 6WCO, 7JU7). Structure 7JU7 obtained in complex with masitinib is one of those outliers (1.84 Å of the first cluster centroid) and was the only one on which the molecule could be redocked. With Smina, up to 8 other molecules could be redocked successfully. Thus, it was retained for the docking. However, to cover a larger number of successfully redocked molecules, we searched for other structures, which when combined provided a larger diversity in the union of successful redocked molecules. We chose the weighting scheme taking chemical diversity into account that is given in the **Supplemental Table 2**.

The best combination with one other structure (5R7Z in addition to 7JU8) gave a score of 8 (max score 29) and successful docking for 12 molecules. The best combination of 3 structures gave a score of 12.1, but only 16 ligands so we kept the second combination (7JU7, 5R7Z, 5RH8), with a score of 11.4, but 18 ligands. It also comprised 5RH8, which gave the best results with FlexX, allowing to diversify the screening with FlexX, while providing a basis for comparison between the two programs.

### Chemical compounds

**GC376**

**MolPort-046-767-490**

**Name**: sodium (2S)-2-[(2S)-2-{[(benzyloxy)carbonyl]amino}-4-methylpentanamido]-1-hydroxy-3-(2-oxopyrrolidin-3-yl)propane-1-sulfonate

**Supplier** : Sigma-Aldrich

**Catalog number** : AMBH93D5391D

**Purity** : ≥ 98%

**Calpain inhibitor XII**

**MolPort-044-183-259**

**Name** : [3-methyl-1-[[[1-[oxo[(2-pyridinylmethyl)amino]acetyl]butyl]amino]carbonyl]butyl]-carbamic acid, phenylmethyl ester

**Supplier** : Cayman Chemical

**Cat number** : 14466

**Purity**: ≥ 90%

**Boceprevir**

**MolPort-046-767-490**

**Name**: sodium (2S)-2-[(2S)-2-{[(benzyloxy)carbonyl]amino}-4-methylpentanamido]-1-hydroxy-3-(2-oxopyrrolidin-3-yl)propane-1-sulfonate

**Supplier**: Sigma-Aldrich; CAS 1416992-39-6) (ref AMBH9614F91F chez Ambeed, Inc) : purity 98%

**Catalog number** : ADV465749229

**Purity**: ≥ 98%

**Mpro 13b**

**MolPort-047-154-487**

**Name:** tert-butyl N-{1-[(1S)-1-{[(2S)-1-(benzylcarbamoyl)-1-oxo-3-[(3S)-2-oxopyrrolidin-3-yl]propan-2-yl]carbamoyl}-2-cyclopropylethyl]-2-oxo-1,2-dihydropyridin-3-yl}carbamate

**Supplier**: Bio-Techne; CAS 2412965-59-2)

**Catalog number**: 7228

**Purity** : ≥ 97% (HPLC)

**Merimepodib**

**MolPort-006-170-006**

**Name**: (3S)-oxolan-3-yl N-{[3-({[3-methoxy-4-(1,3-oxazol-5-yl)phenyl]carbamoyl}amino)phenyl]methyl}carbamate

**Supplier**: Chemscene

**Catalog number**: CS-2105

**Purity**: ≥ 98 %

**Pacritinib**

**MolPort-035-395-848**

**Name**: (16E)-11-[2-(pyrrolidin-1-yl)ethoxy]-14,19-dioxa-5,7,27-triazatetracyclo[19.3.1.1^2^,^6^.1^8^,^12^]heptacosa-1(25),2(27),3,5,8,10,12(26),16,21,23-decaene

**Supplier**: TargetMol

**Catalog number**: T6020

**Purity**: ≥ 99 %

**Fr-PPIchem-27-O18_08616 PAST-0013000**

**MolPort-005-831-261**

**Name**: N-(1,3-benzothiazol-2-yl)-N-(3-chlorophenyl)-4-(1H-imidazol-1-yl)-3-nitrobenzamide

**Supplier**: ENAMINE Ltd.

**Catalog number**: Z27844948

**Purity**: >90%

**Fr-PPIchem-20-D4_06142 PAST-0010527**

**MolPort-016-913-211**

**Name**: 3-{N-[(furan-2-yl)methyl]-1-[2-(phenylamino)-1,3-thiazol-4-yl]formamido}-N-{[3-

(trifluoromethyl)phenyl]methyl}propanamide

**Supplier**: ChemDiv, Inc.

**Catalog number**: V002-9269

**Purity**: >90%

**Fr-PPIchem-08-H7_02385 PAST-0006771**

**MolPort-001-846-784**

**Name**: N-(3,4-dichlorophenyl)-2-[1-(3-methyl-1-benzofuran-2-carbonyl)-3-oxo-1,2,3,4-tetrahydroquinoxalin-2-yl]acetamide

**Supplier**: Vitas-M Laboratory, Ltd.

**Catalog number**: STK823646

**Purity**: >90%

**Fr-PPIchem-33-E11_10265**

**MolPort-006-669-437**

**Name**: 3-[(5-{[(1S,4S,5S)-2-methyl-5-(propan-2-yl)-4-{[4-(pyrimidin-2-yl)piperazin-1-yl]methyl}cyclohex-2-en-1-yl]methyl}-1,3,4-oxadiazol-2-yl)methyl]-1H-indole

**Supplier**: AnalytiCon Discovery, GmbH

**Catalog number**: NAT28-411615

**Purity**: 87.00

**Fr-PPIchem-08-N15_02513 PAST-0006899**

**MolPort-001-583-421**

**Name**: 1-(3-chlorophenyl)-3-(4-{[(3-chlorophenyl)carbamoyl]amino}-1,2,5-oxadiazol-3-yl)urea

**Supplier**: Vitas-M Laboratory, Ltd.

**Catalog number**: STL063220

**Purity**: >90%

